# The reasonable effectiveness of domain adaptation for inference of introgression

**DOI:** 10.1101/2025.01.17.633659

**Authors:** Kerry A. Cobb, Megan L. Smith

## Abstract

Supervised machine learning approaches have proven powerful in population genetics. To use such approaches, training data with known inputs and outputs are required. Since such data are generally unavailable in population genetics, researchers typically rely on simulations under the models of interest to train machine learning algorithms. While powerful, this approach depends heavily on the models used to generate training data. Because of the variety and complexity of processes shaping genetic variation, it is inevitable that not all processes important in an empirical system will be included when generating training data. This leads to a mismatch between the data used to train a machine learning algorithm and the data to which the trained model is ultimately applied–i.e., a domain shift– and can negatively impact inference. Here, we train a Convolutional Neural Network (CNN) to detect introgression between sister populations and demonstrate that it has near perfect accuracy when applied to data generated under the models used for training. To evaluate the impacts of domain shifts on inference, we generated new data with introgression from a third, unsampled population into one of the two focal populations (i.e., ghost introgression), and accuracy was substantially reduced on these data. Finally, we used domain adaptation, which aims to train a network that performs well in the presence of a domain shift. Notably, this requires no knowledge of the target or empirical domain. Our domain adaptation network was able to accurately detect introgression, even in the presence of unmodelled ghost introgression. We also applied this approach to empirical data to detect introgression between ABC Island brown bears and other populations of brown bears. Previous work has suggested that ghost introgression between ABC Island bears and polar bears can mislead tests of introgression between populations of brown bears. We found that using domain adaptation reduced support for introgression between geographically isolated populations of brown bears, suggesting that our approach reduces false inferences of introgression due to ghost introgression.

## Introduction

Analyses of genomic data from across the tree of life have supported rampant introgression between populations and species (Mallet et al., 2016; Taylor and Larson, 2019; Dagilis et al., 2021). Detecting introgression, or the movement of genetic material from one lineage into another, has long interested biologists, as it plays a central role in understanding speciation (Sousa and Hey, 2013; Roux et al., 2016; Payseur and Rieseberg, 2016), inferring species boundaries (Jackson et al., 2017; Smith and Carstens, 2020), developing null models for detecting selection (Nielsen et al., 2007; Excoffier et al., 2009; Luqman et al., 2021), distentangling the factors driving diversification (Thome and Carstens, 2016; Myers et al., 2020; Smith et al., 2022), and more. Given this widespread interest, it is unsurprising that several methods exist for detecting introgression between populations and species. Phylogenetic approaches require data from more than two populations and can only detect introgression between non-sister taxa. Many phylogenetic approaches (e.g., ABBA-BABA tests, Green et al. 2010; *D*_*FOIL*_, Pease and Hahn 2015) evaluate whether there are deviations from the site patterns expected under models without introgression (e.g., the multispecies coalescent model), and interpret deviations from these expected patterns as evidence of potential introgression (reviewed in Hibbins and Hahn 2022). Other phylogenetic approaches aim to infer species networks by quantifying the fit of sequence data to various network topologies under a specific model, such as the multispecies coalescent model on networks (Solís-Lemus and Ané, 2016; Kong et al., 2025). Detecting introgression between sister taxa, on the other hand, requires population genetic approaches and multiple alleles per taxon. Population genetic methods for detecting introgression typically fit a model to the data (e.g., Gutenkunst et al. 2009; Excoffier et al. 2021; Flouri et al. 2020, 2023). Then, migration rates can be inferred and interpreted, or models with and without migration can be compared.

All of these methods rely on models. Phylogenetic approaches rely on models either indirectly (i.e., by comparing expectations under a null model to the observed data) or directly (i.e., by assessing the fit of models to the data), while the most popular population genetic approaches (e.g., Excoffier et al. 2021; Flouri et al. 2023) do the latter. Models must make assumptions about the processes shaping genetic variation. For example, most methods rely on models that assume an absence of gene flow from un-sampled populations (i.e., ghost introgression, Beerli 2004) or that selection has no impact on observed genetic variation. While simplifying assumptions are inevitable in model-based inference, some assumptions are frequently violated in empirical systems, and violations of these assumptions can mislead inference. For example, ghost introgression is almost certainly common in empirical studies. Only a fraction of earth’s biodiversity has likely been described (Costello et al., 2013), and most species that have existed are extinct (Raup, 1992). Even in the absence of extinction and undescribed species-level diversity, empirical studies may fail to sample all relevant populations or species either due to insufficient funds, difficulty sampling, or a lack of knowledge regarding relevant populations. Furthermore, ghost introgression has been shown to mislead both phylogenetic (Tricou et al., 2022) and population genetic (Beerli, 2004; Strasburg and Rieseberg, 2010) approaches for detecting introgression. Similarly, selection appears to influence patterns of variation across the genomes of many species (Cutter and Payseur, 2013; Kern and Hahn, 2018) and can mislead population genetic approaches for detecting introgression (Smith and Hahn, 2024).

Machine learning approaches offer a powerful alternative to maximum likelihood and Bayesian inference when researchers aim to make inferences from highly heterogeneous datasets generated by diverse processes. Specifically, when the models of interest are intractable in likelihood and Bayesian frameworks, supervised machine learning offers a promising solution (Schrider and Kern, 2018; Flagel et al., 2019; Korfmann et al., 2023). Supervised machine learning uses training data—i.e., data with known inputs and outputs—to learn to predict input from output. For example, if a researcher wishes to train a machine learning model to detect introgression, a training dataset consisting of genetic data from pairs of populations or species for which we know whether introgression has occurred is necessary. Of course, such datasets are generally unavailable, especially in the quantities necessary for supervised machine learning (i.e., 1000s of datasets under each model). To circumvent this issue, machine learning approaches in population genetics have typically relied on simulations under the model or models of interest to obtain training data (Schrider and Kern, 2018). This offers a high level of flexibility, as the user can include any processes under which they can simulate data when generating training datasets. Simulations can be used to generate data under a wide array of conditions in a way that is not possible if using very limited empirical data for training, and the hope is that the variation encompassed by the simulations encompasses the empirical data to which a network will later be applied. Supervised machine learning has seen a wide range of successful applications in population genetics, including detecting introgression (e.g., Schrider et al. 2018; Ray et al. 2024; Flagel et al. 2019; Gower et al. 2021), delimiting populations or species (Smith and Carstens, 2020), inferring population size trajectories (e.g., Sanchez et al. 2021), detecting selection (e.g., Hejase et al. 2022), and jointly inferring demography and selection (e.g., Sheehan and Song 2016).

Despite this flexibility, it remains infeasible to simulate every potential process acting in real biological systems. This leads to an inevitable mismatch between the simulated data used to train the machine learning model and the empirical data on which the researcher aims to make predictions. In machine learning, model performance often degrades when models are applied to so-called ‘out-of-distribution’ data, or data that is drawn from a different distribution than the training data. More specifically, a shift between the data used for training and testing is referred to as a domain shift and is known to lead to decreased accuracy. In population genetics and phylogenetics, studies evaluating the impacts of domain shifts on inference have found mixed results. For example, Sheehan and Song (2016) found that applying a network to datasets generated under higher rates of recombination than used in training led to decreased accuracy when detecting selection but did not substantially impact inferences of population sizes. On the other hand, applying a network to datasets generated under more extreme bottlenecks negatively impacted population size inference but had minimal impacts when detecting selection. In another study, S/HIC (Schrider and Kern, 2016), a machine learning approach for detecting hard and soft sweeps, was less impacted than other approaches by misspecified demographic histories. Similarly, Schrider et al. (2018) found minimal impacts of a mis-specified demographic model on attempts to detect introgression. RELERNN (Adrion et al., 2020) estimates recombination rates and also appears to be robust to several model violations. On the other hand, disperseNN (Smith et al., 2023), a neural network for estimating dispersal distance, was negatively impacted when the priors placed on various parameters did not include true values. Thus, the susceptibility of machine learning approaches to domain shifts is likely case-dependent, but is certainly not something that should be ignored.

While domain shifts are potentially a major issue in supervised machine learning, there are approaches that aim to minimize the impacts of such shifts on inference. Domain adaptation approaches aim to construct classifiers that perform well even in the presence of a shift between the source (i.e., training) and target (i.e., empirical) domains (reviewed in Wilson and Cook 2020). Domain mapping and domain-invariant feature extraction are two popular approaches in domain adaptation. Domain mapping aims to map source images to the target domain or vice-versa. Domain mapping often employs an adversarial approach, in which a network is trained to generate target-like images from source images, with similarity assessed using a network trained to discriminate between target and source images (e.g., CycleGAN, Zhu et al. 2017; CyCADA, Hoffman et al. 2018). Here, we will focus on domain-invariant feature extraction, which aims to minimize the differences in features extracted from the source domain and the target domain. Generally, in supervised ML, the aim when extracting features (i.e., the embeddings of data obtained using the learned parameters of the network) is to minimize prediction error (i.e., to find features that maximize prediction accuracy on the training data). In domain-invariant feature extraction, a second optimization goal is added: to find features that are similar between training and target data (e.g., Ganin et al. 2016). Notably, this approach does not require any labeled data (i.e., data with a known history) from the target domain and is therefore potentially powerful in population genetic settings. Mo and Siepel (2023) demonstrated that both background selection and misspecification of the demographic model mislead machine learning approaches for classifying selective sweeps and estimating population recombination rates. However, by using domain-invariant feature extraction, they were able to arrive at accurate inferences in the presence of these processes, even though they were not modeled in the training data. This suggests that domain adaptation may be a promising path forward in population genetic inference.

Here, we first construct a Convolutional Neural Network (CNN) to detect introgression between sister populations. We then evaluate the impacts of a common model violation–ghost introgression–on inferences using this network. Next, we apply a domain adaptation approach to build a new network that performs well in the presence of ghost introgression. Finally, we apply this network to empirical data from brown bears and demonstrate that it reduces the rate of likely false positives.

## Methods

### General Approach

In a general supervised machine learning setting, labeled training data (i.e., **source data**) are used to train a machine learning algorithm. In the case of detecting introgression, labels indicate whether or not introgression has occurred between the focal populations. These data are used as input into a machine learning algorithm, which extracts meaningful features and uses these features to classify data as having been collected from populations that have been completely isolated or have experienced introgression in a binary classification setup. After a network is trained, it can be applied to **target data**. Target data are usually empirical data without known labels, and the goal is to predict a label for the target data. When the models used to generate source data do not capture the processes shaping variation in the target data, a **domain shift** has occurred. **Domain adaptation** techniques aim to train classifiers that work well in the target domain in the presence of a domain shift between the source and target domains (Wilson and Cook, 2020).

Here, we used adversarial domain adaptation, specifically a Conditional Adversarial Domain Adaptation Network (CDAN, Long et al. 2018), which builds on the Domain-Adversarial Neural Network (DANN, Ganin et al. 2016). A traditional CNN performing a classification task has two components: a feature extractor *φ* that maps input data to a feature representation and a classifier *F* that predicts labels from these features. Adversarial domain adaptation adds a third component—a discriminator *D* that attempts to predict whether features came from the source or target domains (Fig. 1). A gradient reversal layer (GRL) between the *φ* and *D* flips the sign of the gradient during backpropagation: *D* is trained to tell domains apart, while the GRL causes *φ* to be trained in the opposite direction (i.e., to produce features that *D* cannot use to distinguish domains). Given labeled source data (*X*_*S*_, *y*_*S*_) and unlabeled target data (*X*_*T*_), DANN optimizes

**Figure 1.**
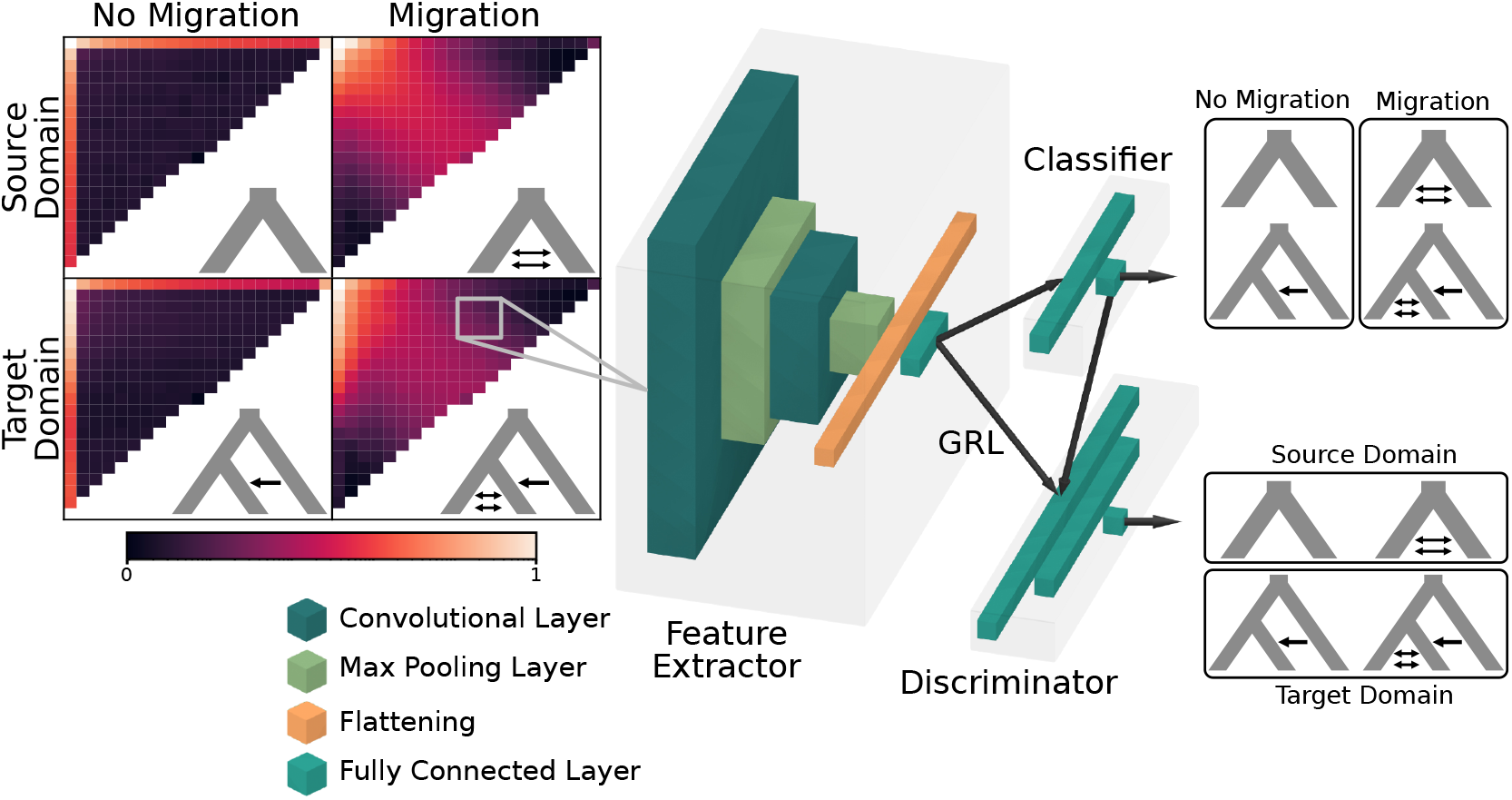
Overview of our CDAN architecture. Genetic data from a pair of populations are summarized as joint Site Frequency Spectra (SFS) which are used as input into the network. The feature extractor *φ* extracts features from the SFS. The classifier *F* uses these features to detect introgression, while the discriminator *D* uses these features along with the classifier predictions to distinguish the source and target domains. However, the gradient reversal layer (GRL) inverts the loss function during back propagation, which discourages the feature extractor from learning features that are not shared by both domains, minimizing the domain discrimination.

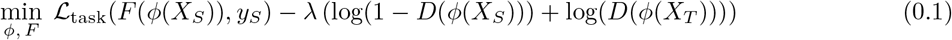

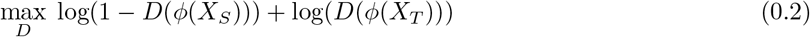

where *λ* sets the trade-off between task performance and domain confusion. Because of the GRL, minimizing Eq. (0.1) and maximizing Eq. (0.2) causes *D* to become better at distinguishing domains while *φ* learns to make that more difficult, ideally yielding domain-invariant features.

CDAN (Long et al., 2018)) extends DANN by conditioning the discriminator on *F* ‘s predictions, which better captures cases in which the joint distributions of features and classes differs across domains. Rather than concatenating the features and predictions, CDAN uses the multilinear map of the features and the predictions (*f* ⊗ *g*), which captures the cross-covariance between the two and has been shown to outperform DANN (Long et al., 2018). The objective becomes:

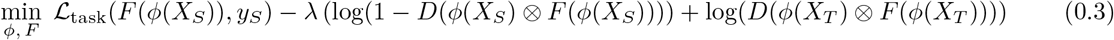

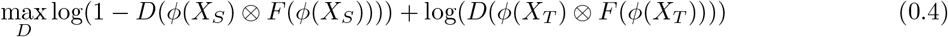

This is a slight simplification of the actual loss function used here, as we also used entropy regularization, which prioritizes easy-to-transfer instances. See the documentation of adapt (de Mathelin et al., 2021) for the exact loss functions implemented. Below, we describe our approach for simulating training and target data, the details of our network, and an empirical test using data from brown bears.

### Simulations

To generate data for training and testing our machine learning approach for detecting introgression, we used msprime v1.3.1 (Baumdicker et al., 2022). For our source data, we focused on two demographic models: a divergence-only model and a model with gene flow upon secondary contact between the two focal populations (Fig. 2) We chose these models because detecting gene flow between sister populations is a commonly undertaken task in evolutionary biology, and secondary contact models are often considered, for example in regions impacted by Pleistocene glacial cycles (e.g., Smith et al. 2022). We simulated a 500,000 bp region of a single chromosome with a mutation rate *µ* and a recombination rate *r* (Table 1). The populations diverged at time *τ* and shared a single constant effective population size. For the secondary contact model, migration began halfway between the time of divergence and the present and continued until the present. Population sizes, divergence times (*τ*), and migration rates were drawn from uniform priors (Table 1). We sampled 10 diploid individuals from each population at the present and summarized the data as folded joint site frequency spectra (SFS). We then normalized each SFS output from msprime such that all values sum to one.

**Table 1.** Parameter values and ranges of uniform probability distributions for simulations in msprime. All times are in units of generations. Mutation and recombination rates are in units of per site per generation. The migration rate is the effective population size *N*_*e*_ times the expected number of migrants per generation *m*. The pulse migration proportion is the proportion of lineages that move from one population into another at time *τ*_*pulse*_ looking backward in time.

|  | General Scenario | Bear Scenario |
| --- | --- | --- |
| Haploid sample number | 20 | 20 |
| Sequence length | 500,000 | 10,000,000 |
| Mutation rate | $1e-8$ | $1e-8$ |
| Recombination rate | $1e-8$ | $1e-8$ |
| Effective population size ($N_e$) | 10,000–100,000 | 1,000–20,000 |
| Divergence time ($\tau$) | 50,000–500,000 | 1,000–10,000 |
| Secondary contact time ($\tau_{migration}$) | $\tau/2$ | $\tau/2$ |
| Migration rate ($N_e * m$) | 0.05–5.0 | 0.05–5.0 |
| Ghost divergence time ($\tau_{ghost}$) | 1,000,000–2,000,000 | 20,000–100,000 |
| Ghost pulse migration time ($\tau_{pulse}$) | 500–2,000 | $0-0.75 * \tau$ |
| Ghost pulse migration proportion ($\gamma$) | 0.05–0.3 | 0.05–0.3 |

**Figure 2.**
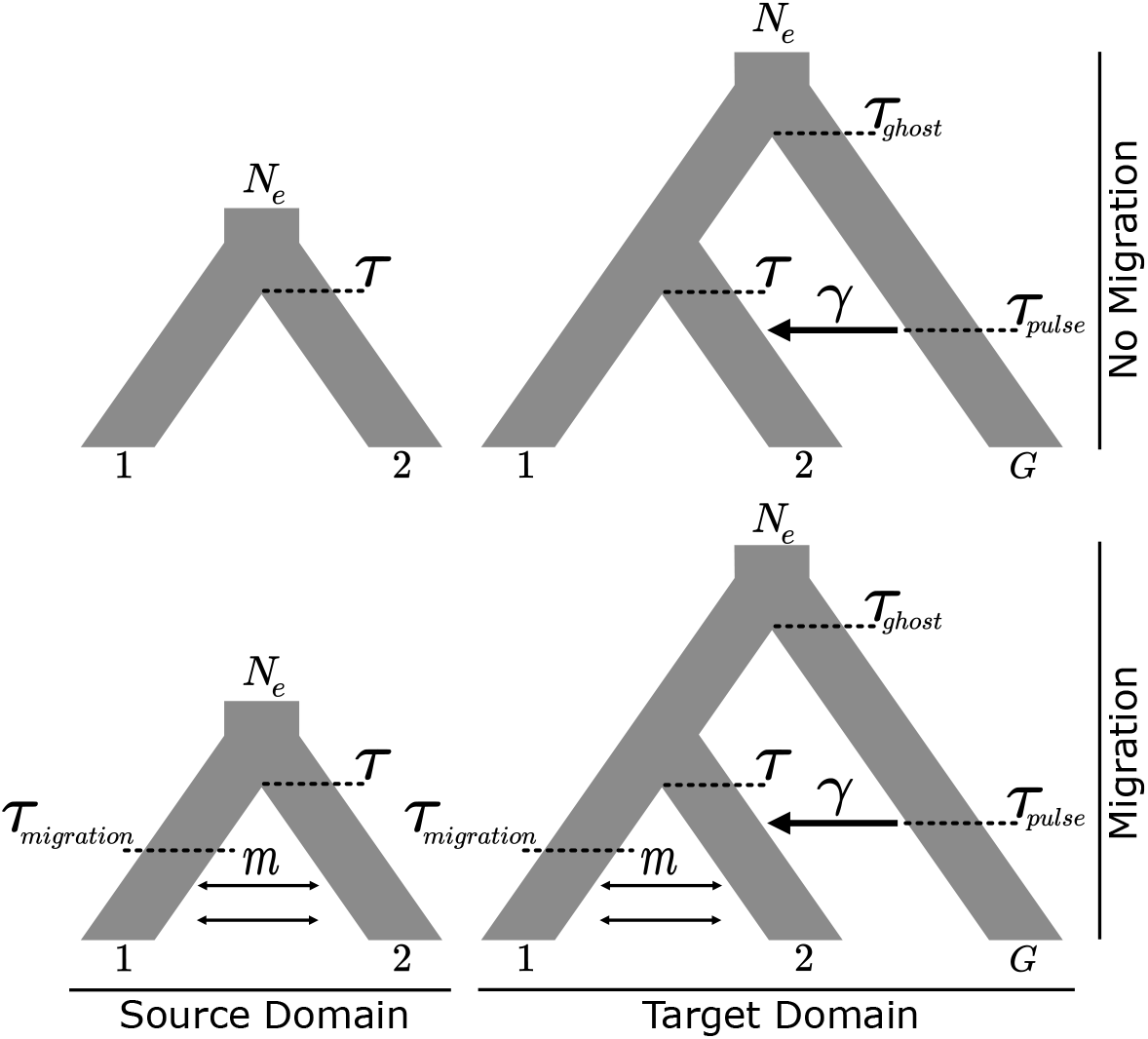
Models used to simulate training and testing data. In the source domain, we model two taxa with or without migration upon secondary contact. In the target domain, we expand upon these models by adding introgression into taxon 2 from unsampled taxon *G*–a ghost lineage. The effective population size *N*_*e*_ is shared among all populations. Divergence between taxa 1 and 2 occurs at time *τ*, and divergence between the ancestor of these taxa and taxon *G* occurs at time *τ*_*ghost*_. When migration occurs, it begins at time *τ*_*migration*_ and continues until the present at a rate *m*. Introgression from the ghost population is modeled as a pulse event in which a proportion *γ* of lineages move from taxon 2 to taxon *G* (backwards in time) at time *τ*_*pulse*_.

Next, we simulated data under an alternative model which we treat as the target domain. Data from the target domain are typically unlabeled. However, it is useful to have target data with known labels to evaluate the impact of mis-specification on a standard machine learning approach and the effectiveness of domain adaptation. While known, these labels are not used for inference. Specifically, we simulated data under a model with introgression from an un-sampled “ghost” population into one of the two focal populations. Our ghost introgression models were identical to the two models described above, except they also included a third unsampled population that diverged from the two focal populations at time *τ*_*ghost*_. Looking backward in time, at time *τ*_*pulse*_, a proportion of lineages *γ* from one of the focal populations moved into the ghost lineage (Fig. 2). The *γ, τ*_*ghost*_, and *τ*_*pulse*_ parameters were drawn from uniform priors (Table 1). All other aspects of the models were parameterized as in the source models. Together with the source model parameters, we refer to these parameters as the “general scenario”. As above, we summarized the simulated data as a normalized, folded joint SFS for the two focal populations with the recipient of ghost introgression on the row axis. o visualize the impacts of migration and ghost introgression on the SFS, we plotted SFS and Principal Component Analyses (PCAs) of the SFS under the source and target domains, with and without migration.

### Network

We used these simulated data as input for training and testing a CDAN. We implemented our CDAN using the python package adapt v0.4.3 (de Mathelin et al., 2021), along with functions from tensorflow v2.11.1 (Abadi et al., 2015) using the keras API (Chollet et al., 2015). For the feature extractor *φ*, we used two successive blocks containing a 2D convolutional layer with a 3 × 3 filter, a 1 × 1 stride, and ReLU activation followed by 2D max pooling with a 2 × 2 pool size and a 2 × 2 stride. These blocks were followed by flattening and a dense layer with 20 neurons and ReLU activation. For the classifier *F*, we used a dense layer with 100 neurons and ReLU activation followed by a dense layer with 2 neurons and softmax activation. For the discriminator *D*, we used a dense layer with 100 neurons followed by another with 50 neurons, each with ReLU activation and, finally, a dense layer with a single neuron and a sigmoid activation function. We used a categorical cross-entropy loss function for the classifier. We used the Adam optimizer with a learning rate of 0.001 for each component of the network. The CDAN parameter *λ* is a trade-off parameter that controls the relative weight given to correctly classifying the source data versus minimizing differences between the source and target features. We used a variable *λ* which was initially set to zero and was then increased linearly up to its specified value over the first 1000 mini-batches across epochs. As is common in machine learning applications, our network includes many hyperparameters. During initial tests of this approach, we explored a range of batch sizes, learning rates for the feature extractor, classifier, and discriminator, and values of the tradeoff parameter *λ*. Given the high performance obtained using the values described above, we did not deem more hyperparameter tuning necessary for this application.

### Training and Evaluation

We used data simulated under the models described above to train and evaluate our networks. Under the source domain (i.e., with no ghost introgression), we simulated 20,000 training datasets, 1,000 test datasets, and 1,000 validation datasets. In analyses of empirical data, we expect to have far less data from the target domain, as this domain will be comprised entirely of empirical data. Therefore, to evaluate domain adaptation under a realistic scenario, we simulated only 100 datasets for training under the target domain, along with 1,000 test datasets for assessing performance. In order to compare networks trained without domain adaptation to networks trained with domain adaptation, we varied the value of the trade-off parameter *λ*. When *λ* is set to zero, it is equivalent to using only the feature extractor and classifier, thus having no domain adaptation. We trained 10 replicate networks with *λ* set to zero and evaluated the performance of these networks on test data generated under both the source and target domain. Then, we trained 10 replicate networks with a maximum *λ* of 1 to evaluate the effectiveness of domain adaptation and evaluated these networks on the same test datasets. All networks were trained for 50 epochs using a batch size of 16. Network performance was evaluated using confusion matrices and receiver operator curves (ROC). Additionally, we performed principal components analysis (PCA) of feature embeddings for all test datasets. Feature embeddings are low-dimensional representations of high-dimensional inputs. Here, they are the outputs obtained by passing the input SFS through the feature extractor branch *φ* of the network. Visualizing these embeddings allows us to assess differences in the features extracted from the source and target domains. If a domain shift has occurred, we expect to see differences between these distributions. However, if domain adaptation is successful, such differences should be diminished.

### Performance On Imbalanced Target Data And Importance Weighting

Above, both source and target data were composed of equal numbers of observations from the two classes (migration and no migration). However, in empirical datasets, the assumption of balanced classes is unlikely to be met. In the machine learning literature, this is referred to as a ‘label shift’, and label shifts have been shown to negatively impact the performance of adversarial domain adaptation approaches (Zhao et al., 2019). To assess the impacts of class imbalance, we subsampled our target data, keeping 50 total samples, and we varied the proportion of samples from the no migration class across five levels: 0, 0.25, 0.50, 0.75, and 1.0. We then trained CDAN classifiers as described above, and assessed overall accuracy on the subset of target data used for training.

Given the striking decrease in performance in the presence of imbalance (see Results), we explored the use of importance weights to improve the robustness of our classifiers to label shifts (Tachet des Combes et al., 2020). We implemented an importance-weighted version of CDAN (IWCDAN) based on the architecture described by des Combes et al. (2020) to account for label shift during domain adaptation. Briefly, IWCDAN weights the adversarial discriminator loss on the source data during training based on the estimated prevalence of predicted classes in the target data. During training, we estimated class importance weights from the source confusion matrix and the predicted class distribution for the unlabeled target data. Using these weights, we reweighted the source component of the adversarial loss. Importance weights were updated every five epochs. To improve stability, we updated importance weights using an exponential moving average with a momentum of 0.5. Initially, we observed that IWCDAN was very sensitive to *λ*, so we explored four values of *λ*: 0, 0.1, 0.5, and 1.0. We did not change *λ* across epochs as above, as initial runs indicated that this introduced additional instability.

### Comparison To Fastsimcoal2

To evaluate whether a more widely used approach that relies on the SFS is vulnerable to ghost introgression, we analyzed 100 datasets each for the divergence and secondary contact models, with and without ghost introgression, in fastsimcoal2 v2.8.0.0 (Excoffier et al., 2021). To reduce the impacts of linkage, we subsampled our data, only including SNPs that were separated by at least 500 bp. We compared two models–a divergence-only model and a secondary-contact model, with priors set identical to those used to simulate the data. We used 100,000 simulations to estimate the composite likelihood and 40 cycles of the Brent algorithm for parameter estimation. We used the folded SFS and ignored monomorphic sites. For each simulated dataset, we performed ten replicates and selected the replicate with the highest likelihood. We then performed a Likelihood Ratio test and rejected the divergence-only model at an *α* level of 0.01.

### Application To Brown Bears

Brown bears are widely distributed across the Northern hemisphere and exhibit a high degree of population structure (de Jong et al., 2023). Beginning around the time of the last glacial maximum, several distinct populations diverged (de Jong et al., 2023). Some of these populations have experienced introgression from polar bears, with the most well-known case being brown bears from the Admiralty, Baranof, and Chichagof (ABC) islands in Southeastern Alaska (Cahill et al., 2015; Kumar et al., 2017; Wang et al., 2022; Lan et al., 2022). Introgression between adjacent but divergent brown bear populations is plausible, whereas introgression between non-adjacent populations is unlikely. However, introgression has been inferred between very distant populations when ghost introgression from the polar bear lineage is not considered, making brown bears a good test case for our approach (Tricou et al., 2022). We simulated source and target data under a scenario similar to the history of polar bears and brown bears (see below for exact parameters) and trained models using the same network architecture as above in order to evaluate the performance of our approach in this context. We then applied our approach to test for introgression between ABC Island bears and seven other brown bear populations using networks trained with and without domain adaptation. In this case, our target data are unlabeled empirical data.

#### Testing under the bear scenario using simulations

First, we used a simulation strategy similar to that above, but with parameters suitable for the bear scenario, to assess the performance of networks with and without domain adaptation under these conditions. To generate simulated data for training, validation, and testing under the “bear scenario”, we used the same simulation procedure as above, changing only the parameters of the models (Table 1). In addition to simulations for training our classifier in the source domain, we simulated data meant to mimic the target domain to evaluate the efficacy of our approach in this setting. In this case, in the target domain we anticipate introgression (forward in time) from the polar bear lineage into the ABC Island bears. Polar bears are estimated to have diverged from brown bears 0.38–1.05 mya. (Kutschera et al., 2014; de Jong et al., 2023; Abella et al., 2012; Zou et al., 2022). This is equivalent to 30,400–84,000 generations assuming a mean generation time of 12.5 years as estimated by Wang et al. (2022). The divergence times between pairs of brown bear populations have been estimated to be between 25,000–80,000 years or 2,000–6,400 generations ago assuming the same mean generation time (de Jong et al., 2023). We drew effective population sizes from a uniform prior spanning 1,000–20,000, which encompasses values used in previous simulation studies (de Jong et al., 2023). We used a mutation rate of 1*e* − 8 which approximates the estimate of 0.84e-8 by Wang et al. (2022). We followed de Jong et al. (2023) in setting the recombination rate to 1*e* − 8. As before, we simulated 20,000 training datasets, 1,000 test datasets and 1,000 validation datasets under the source domain (i.e., with no polar bear introgression), and we simulated 100 training datasets and 1,000 test datasets under the target domain (i.e., with polar bear introgression). We summarized these simulated data as normalized, folded joint SFS. We trained and tested networks using the same procedure used for the general scenario with one minor modification. When training networks with domain adaptation, we used a maximum *λ* of 0.5 because during preliminary analyses we observed that a *λ* of 0.5 resulted in better, more stable training performance relative to a *λ* of 1.0 for the bear scenario. We trained 10 replicate networks for each *λ* value (i.e., *λ*=0 for no domain adaptation and *λ*=0.5 for domain adaptation). We report both average softmax probabilities and binary classifications based on thresholds (e.g., classifying chromosomes as introgressed if the average softmax probability is > 0.5). We evaluated training performance as we did for the general scenario. We also explored the performance of IWCDAN, as described above.

#### Applying the networks to empirical data from brown bears

After finding that IWCDAN worked well in a simulation setting similar to the expected history of brown and polar bears (see Results), we applied importance-weighted networks to empirical target data from brown bear populations. In order to train domain adapted networks with empirical target data and test for introgression between ABC Island bears and the seven other brown bear populations, we used publicly available whole genome re-sequencing data (Fig. 3) (de Jong et al., 2023; Benazzo et al., 2017; Barlow et al., 2018; Cahill et al., 2013, 2015; Liu et al., 2014; Miller et al., 2012; Taylor et al., 2018). For each of the eight populations, we arbitrarily selected 10 samples and obtained all of the available Illumina sequencing run data for each sample from the European Nucleotide Archive (see Table S2 for accession IDs). We used fastp v0.23.4 to remove any remaining adapter sequences and to remove low quality sequences using the default parameters (Chen, 2023). We mapped the filtered and trimmed reads to the 36 largest autosomal scaffolds (assembled chromosomes) of the brown bear genome published by Armstrong et al. (2022) using BWA-MEM2 v2.2.1 (Li, 2013) with default settings. We merged any mapped reads from samples that were split across different runs using Samtools v1.20 (Danecek et al., 2021). To call variants from the mapped reads we used BCFtools v1.20, running the mpileup command with a max depth of 250 and without INDEL calling (Danecek et al., 2021). Nextflow (Di Tommaso et al., 2017) scripts used for processing the bear data are archived at https://doi.org/10.5281/zenodo.14629180. For each of the seven population pairs, we filtered any sites that had missing data and retained only biallelic sites among the samples within the two populations using BCFtools. We then summarized the filtered data from each chromosome as a folded, joint SFS with ABC Island bears on the row axis using *δ*a*δ*i v2.3.3 (Gutenkunst et al., 2009). We treated chromosomes as independent for several reasons: 1) patterns of introgression are expected to vary across the genome, and understanding how introgression varies across chromosomes could be of biological interest; 2) using a more complicated simulation scheme with multiple chromosomes seems largely unnecessary for the task of detecting introgression, as we had high power with only 500,000 bp regions in our simulation study; and 3) we suspect that having semi-independent replicates (due to recombination) may be helpful during domain adaptation. When the classification of an allele as the major or minor allele in one population versus the other is ambiguous, *δ*a*δ*i adds 0.5 to each of the two possible cells in the folded, joint SFS. This differs from msprime where alleles are always classified as the minor allele in the population on the row axis if classification is ambiguous. To make the SFS output by *δ*a*δ*i consistent with the SFS output by msprime, we summed both counts of ambiguously classified alleles and replaced the minor allele count in the population on the row axis with this sum and the major allele count with 0. We then normalized the allele counts as above.

**Figure 3.**
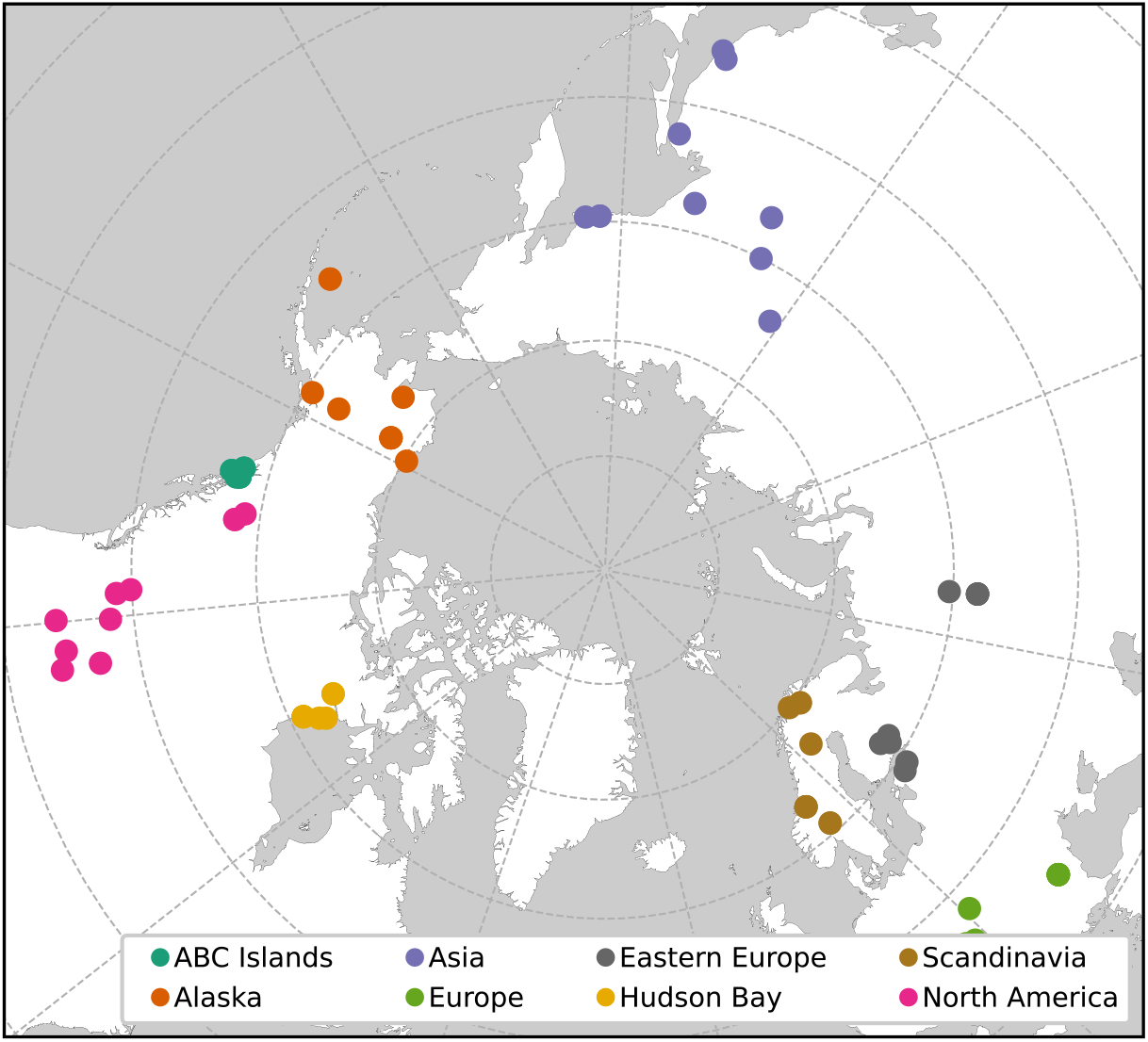
Localities and population assignments for brown bear samples used in this study. We used the North Polar Stereographic projection to map occurrences. Colors indicate the different populations.

We trained 10 replicate networks with a *λ* of zero, 10 replicate networks with a *λ* of 0.1, and 10 replicate networks with a *λ* of 0.5 using IWCDAN as described above, given its superior performance in the presence of class imbalance, which may be present in these data. For the source training data we re-used the 20,000 training and 1,000 validation datasets used above under the bear scenario. For the target training data we used the 36 SFSs computed from each chromosome from all population pairs involving the ABC Island bears. We then computed softmax probabilities of introgression for each chromosome in each population pair independently using the trained network, and we classified chromosomes as introgressed or not using a classification threshold of 0.5.

## Results

### General Scenario Simulation

Both migration and ghost introgression leave clear signals in the simulated datasets. Datasets generated under the source domain had an average of 8,917 Single Nucleotide Polymorphisms (SNPs), and migration between populations led to increased allele sharing in the SFS, whereas, in the absence of migration, there were more alleles unique to each population (Fig. 1, Fig. S1, Fig. S2). Datasets generated under the target domain (i.e., with ghost introgression) contained more SNPs on average (22,032). Furthermore, ghost introgression tended to skew the SFS. Perhaps most notably, ghost introgression led to an increase in the number of alleles unique to the recipient of ghost introgression (population 2, Fig. S3a,b, Fig. S4a,b). In the absence of migration between the focal populations, ghost introgression also decreased the number of fixed alleles in population 1 (Fig. S3a, Fig. S4a). Finally, ghost introgression led to a very slight excess of shared variants between the focal populations, even in the absence of migration, although this pattern is only obvious when observing the unnormalized SFS (Fig. S4c). While ghost introgression does not obviously drive no migration scenarios to look identical to migration scenarios, there are substantive differences between the source and target domains, as also evidenced in the PCA (Fig. S1). Notably, there appears to be more overlap between datasets with and without migration when ghost introgression is present.

Our CNN without domain adaptation (i.e., *λ* = 0) had high power and a near-zero false positive rate in the absence of a domain shift. Introgression was never inferred for datasets lacking introgression, and networks failed to detect introgression when it was present in a mean of 0.14% of datasets across replicate networks (Fig. S5a, Table S1). The mean AUC score was greater than 0.999, indicating a near perfect classifier (Fig. 4b).

**Figure 4.**
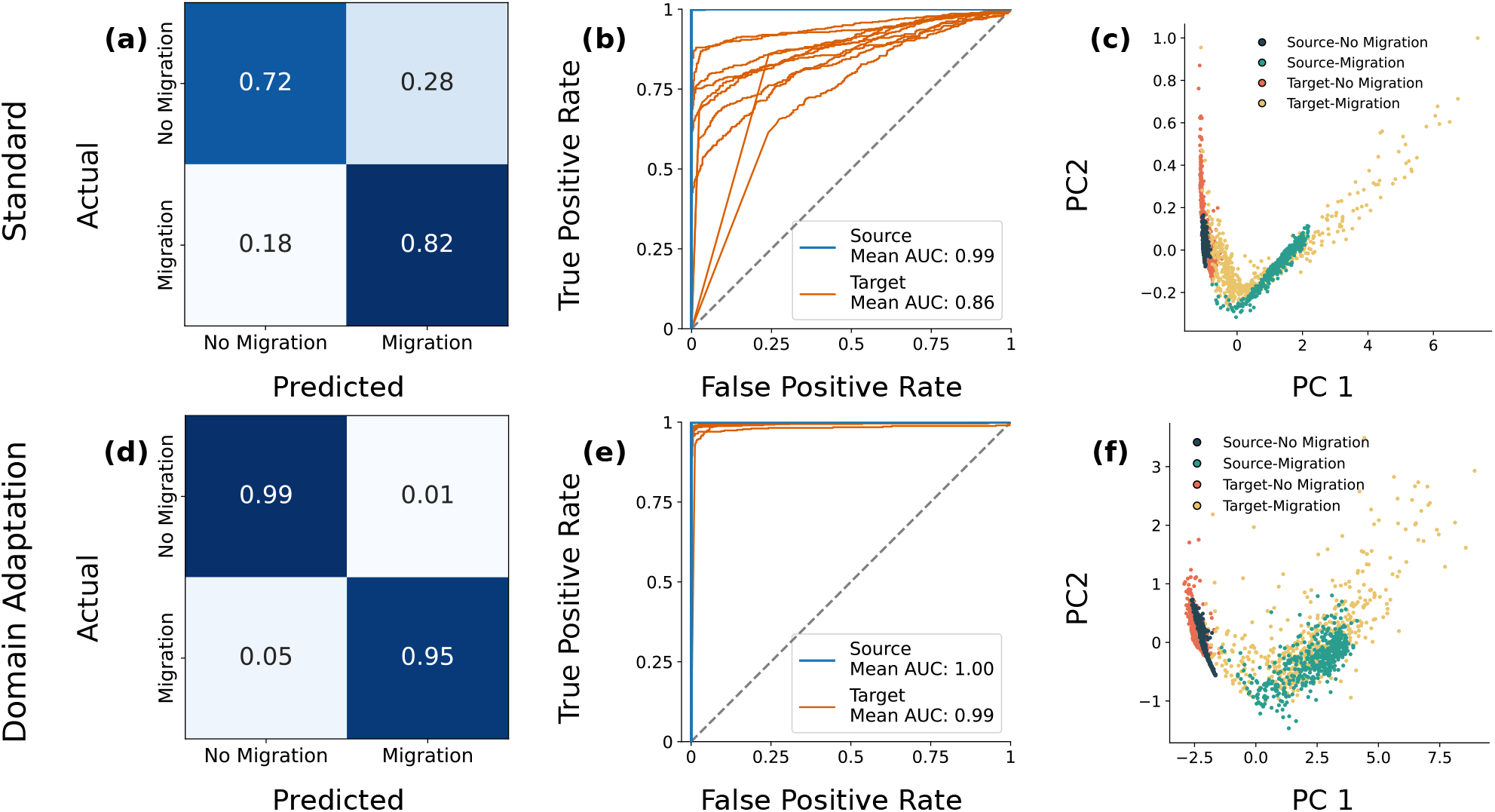
Results under the general simulation scenario on datasets generated under the target domain (i.e., with ghost introgression). Confusion matrices show the proportion of test replicates in each category without (a) and with (d) domain adaptation. Values were averaged across the ten replicate networks. (b) ROC curves for the source (blue) and target (orange) test datasets without (b) and with (e) domain adaptation. The average AUC across all replicates is reported. (c) PCA of feature embeddings from the source (blue) and target (orange) test datasets without (c) and with (f) domain adaptation from an exemplar replicate. All values are rounded to the nearest hundredths place. See Table S1 for original values.

However, in the presence of ghost introgression, the network trained without domain adaptation performed substantially worse. Introgression was incorrectly inferred when it was absent in a mean of 28.10% of datasets, and networks failed to infer introgression when it was present in a mean of 17.98% of datasets (Fig. 4a). The mean AUC score was 0.86 with a high degree of variation observed in the ROC curves (Fig. 4b). Furthermore, the PCA of source and target embeddings indicated a high degree of mismatch between the embedded features of the two domains (Fig. 4c). Notably, datasets generated with ghost introgression tended to have a wider spread than those generated under the source domain, leading to more overlap between datasets generated with and without migration between the focal populations. This is particularly notable in the case of target datasets without migration, which may explain the high false positive rates observed here.

Next, we evaluated the impact of using domain adaptation on inferences with and without simulation misspecification. In the absence of simulation misspecification, using domain adaptation had no substantial negative impact on network performance. There was only a single false positive across all replicates, and networks failed to detect introgression in a mean of 0.3% of datasets (Fig. S5b). The mean AUC score across replicates was greater than 0.999, again indicating a near perfect classifier (Fig. 4e).

In the presence of ghost introgression, using domain adaptation substantially improved accuracy compared to the networks without domain adaptation. Introgression was incorrectly inferred when it was absent in a mean of only 0.86% of datasets across replicates, and networks failed to infer introgression when it was present in a mean of only 5.04% of datasets across replicates (Fig. 4d). The mean AUC score was 0.994, indicating a near perfect classifier even in the presence of simulation misspecification (Fig. 4e). The PCA of source and target embeddings also indicated a greater degree of overlap between the domains than was seen with the network trained without domain adaptation (Fig. 4f).

Classification accuracy increased early in training, and the accuracy remained high throughout (Fig. S6). On the other hand, discriminator accuracy often began high, but decreased quickly, fluctuating around 0.5 for most of training. This indicates that our network performs as desired, extracting features that facilitate classification while minimizing differences between the source and target domains.

### Performance On Imbalanced Target Data and Importance Weighting

In the absence of importance weighting and when *λ >* 0, accuracy on the target data declined substantially with increasing imbalance (Fig. 5). The impact of importance weighting on target accuracy varied strikingly across values of *λ* (Fig. 5), while the impact on source accuracy was negligible (Fig. S7). When *λ* was zero, target accuracy remained low regardless of the proportion of datasets from the target, as expected. However, when *λ* was 0.1, importance weighting improved accuracy relative to the baseline under moderate class imbalance (i.e., when the target proportion was 0.25 or 0.75). Notably, accuracy remains above that seen when *λ* = 0, except in extreme cases of class imbalance (0.0 or 1.0). When all datasets come from a single class, IWCDAN cannot rescue performance, and domain adaptation is unsuccessful, and, in fact, degrades performance compared to the *λ* = 0 case, while at higher values of *λ*, the performance of domain adaptation also degrades. Critically, it is possible to diagnose cases in which *λ* is too high by looking at the importance weights across replicates and epochs (Fig. S8). When *λ* is 0.1, the importance weights quickly converge on reasonable values across all replicates. At higher values of *λ* importance weights are more variable across replicates. Thus, these weights could be used to diagnose issues and select the optimal *λ*.

**Figure 5.**
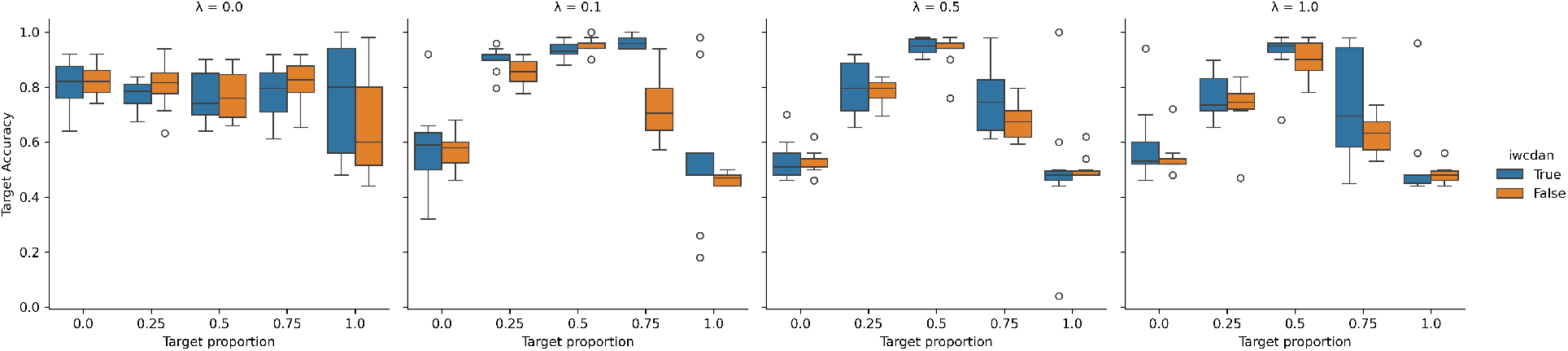
Results across imbalanced datasets under the general simulation scenario under the target domain. The x-axis is the proportion of the 50 target datasets coming from the no migration class. Blue boxplots show results with importance weighting, and orange boxplots show results without importance weighting.

### Comparison To Fastsimcoal2

Fastsimcoal2 was minimally impacted by ghost introgression. In the absence of ghost introgression, fastsimcoal2 correctly classified all secondary-contact datasets and misclassified a single divergence-only dataset (1% overall error rate). When ghost introgression occurred, fastsimcoal2 correctly classified all divergence-only datasets and misclassified two secondary-contact datasets (2% overall error rate). This suggests that fastsimcoal2 is relatively robust to ghost introgression.

### Bear Scenario Simulation

The bear scenario presents a more challenging problem than the general scenario due to more recent divergence times, so we also evaluated the performance of networks with and without domain adaptation under this scenario. The source datasets had a mean of 21,851 SNPs, while target datasets had a mean of 34,112 SNPs. Unsurprisingly, even in the absence of simulation misspecification, networks performed slightly worse in this scenario, but error rates remained low. Networks trained without domain adaptation (i.e., *λ* = 0) falsely inferred introgression when it was absent in a mean of 4.14% of datasets and failed to detect introgression when it was present in a mean of 6.22% of datasets (Fig. S5c). The mean AUC score for datasets without simulation misspecification was 0.991, indicating a highly accurate classifier (Fig. 6b).

**Figure 6.**
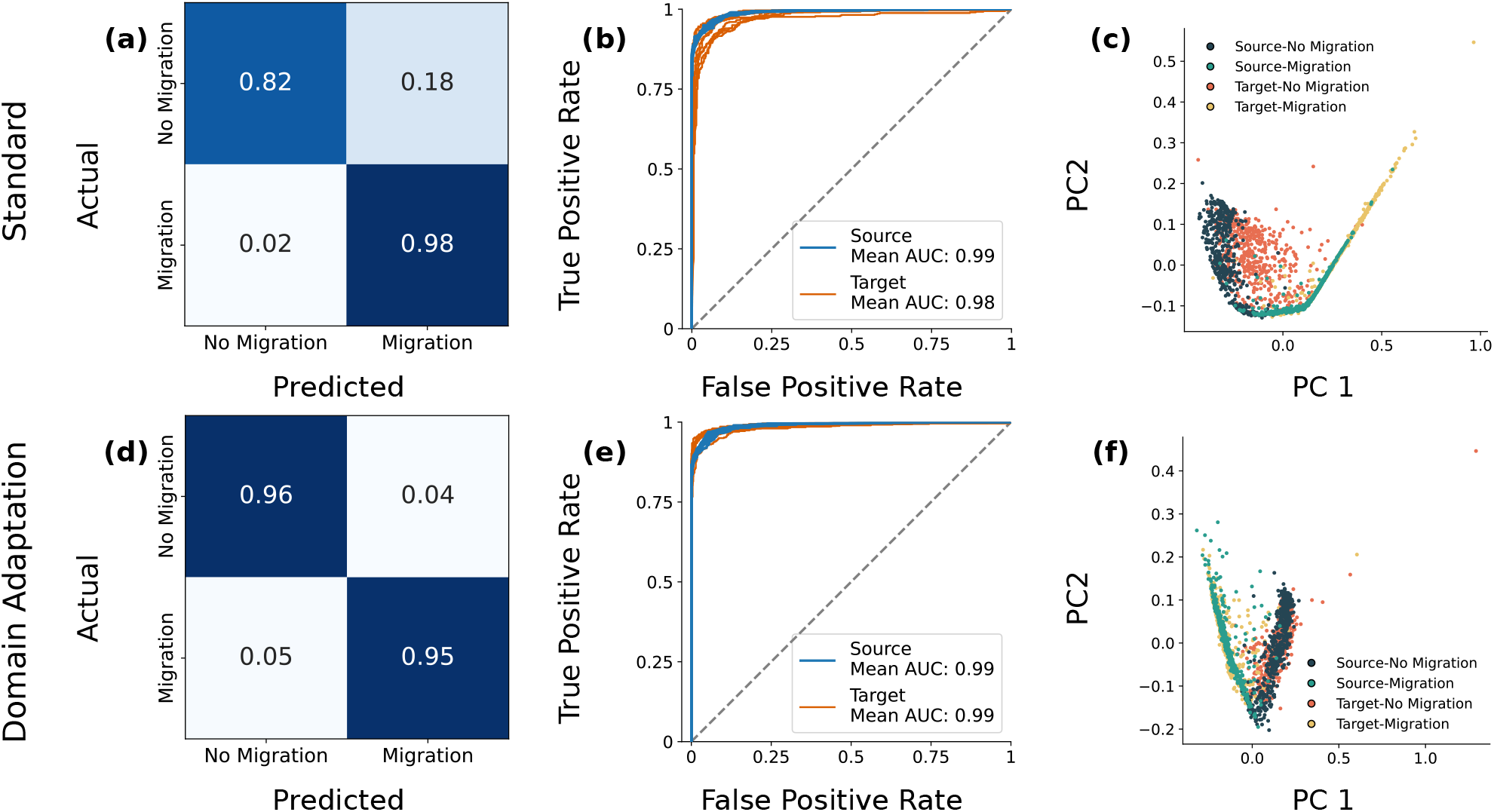
Results under the bear simulation scenario on datasets generated under the target domain (i.e., with ghost introgression). Confusion matrices show the proportion of test replicates in each category without (a) and with (d) domain adaptation. Values were averaged across the ten replicate networks. (b) ROC curves for the source (blue) and target (orange) test datasets without (b) and with (e) domain adaptation. The average AUC across all replicates is reported. (c) PCA of feature embeddings from the source (blue) and target (orange) test datasets without (c) and with (f) domain adaptation from an exemplar replicate. All values are rounded to the nearest hundredths place. See Table S1 for original values.

Network performance was again substantially worse in the presence of ghost introgression. Introgression was incorrectly inferred when it was absent in a mean of 17.6% of datasets across replicates, and networks failed to infer introgression when it was present in a mean of 2.08% of datasets (Fig. 6a). However, the mean AUC score across replicates for tests with ghost introgression was 0.98, which is nearly equal to the mean AUC score for tests using source data, likely due to high performance on the positive class (Fig. 6b). In light of this, we explored the possibility of using a more stringent classification threshold in lieu of domain adaptation. However, although performance on the target class was improved at a threshold of 0.9, it did not match performance using domain adaptation, and false negatives became much more prevalent in the source domain (Fig. S9). The PCA of source and target domain embeddings indicated some degree of mismatch between the two domains, again with greater spread in the target domain (Fig. 6c).

As in the general scenario, domain adaptation did not negatively impact performance in the absence of simulation misspecification. Networks falsely inferred introgression when it was absent in a mean of 4.7% of datasets and failed to detect introgression when it was present in a mean of 5.68% of datasets (Fig. S5d). The mean AUC score across replicates for datasets without simulation misspecification was 0.99 (Fig. 6e).

As in the general scenario, in the presence of ghost introgression, networks with domain adaptation substantially outperformed networks without domain adaptation. The trained networks misclassified a mean of 4.38% of datasets simulated without introgression and 4.82% of datasets simulated with introgression (Fig. 6d). The mean AUC score across replicates was 0.989 (Fig. 6e). The PCA of source and target domain embeddings had a greater degree of overlap than the embeddings from the network without domain adaptation (Fig. 6f).

As in the general simulation scenario, imbalance across classes in the target data led to a substantial drop in performance on the target data when *λ* was greater than zero (Fig. 7), while accuracy on the source data remained high (Fig. S10). Again as above, the impact of importance weighting varied across values of *λ*. When *λ* was 0.1, importance weighting improved accuracy under moderate class imbalance, exceeding accuracy in the absence of domain adaptation (i.e., when *λ* = 0). However, importance weighting could not rescue performance when all samples came from a single class. At higher values of *λ*, importance weighting was, again, less effective. As above, the estimated importance weights could be used to diagnose issues and select the optimal *λ* (Fig. S11).

**Figure 7.**
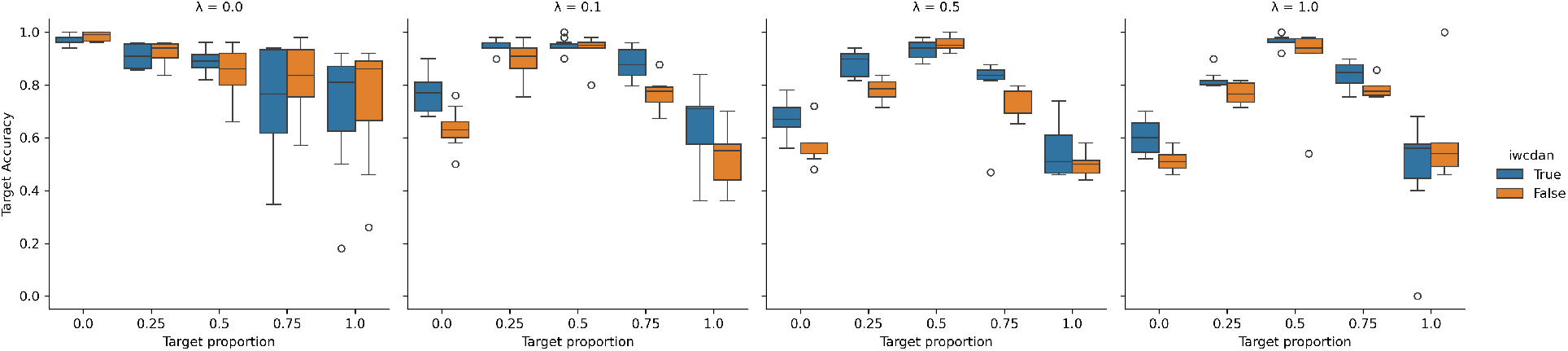
Results across imbalanced datasets under the bear simulation scenario under the target domain. The x-axis is the proportion of the 50 target datasets coming from the no migration class. Blue boxplots show results with importance weighting, and orange boxplots show results without importance weighting.

### Empirical Bear Data

Next, we applied networks with and without domain adaptation to data from the ABC Island bears and seven other brown bear populations, with the expectation that ghost introgression from polar bears to the ABC Island bears could mislead inferences. Given results on simulated data, we always used importance weighting in the empirical analyses. After mapping the bear sequence data to the reference and filtering, the mean number of SNPs per chromosome in each population pair tested for introgression was 497,657.4. Without domain adaptation, the softmax probability of introgression across replicates was high across most chromosomes and population pairs, with a mean of 0.79 (SD = 0.27) (Fig. 8a). Notably, inferred probabilities of introgression were high even between the ABC Island bears and geographically distant populations from Asia and Europe. When applying a classification threshold of 0.5, most chromosomes from most populations were classified as having introgression (Fig. 8d).

**Figure 8.**
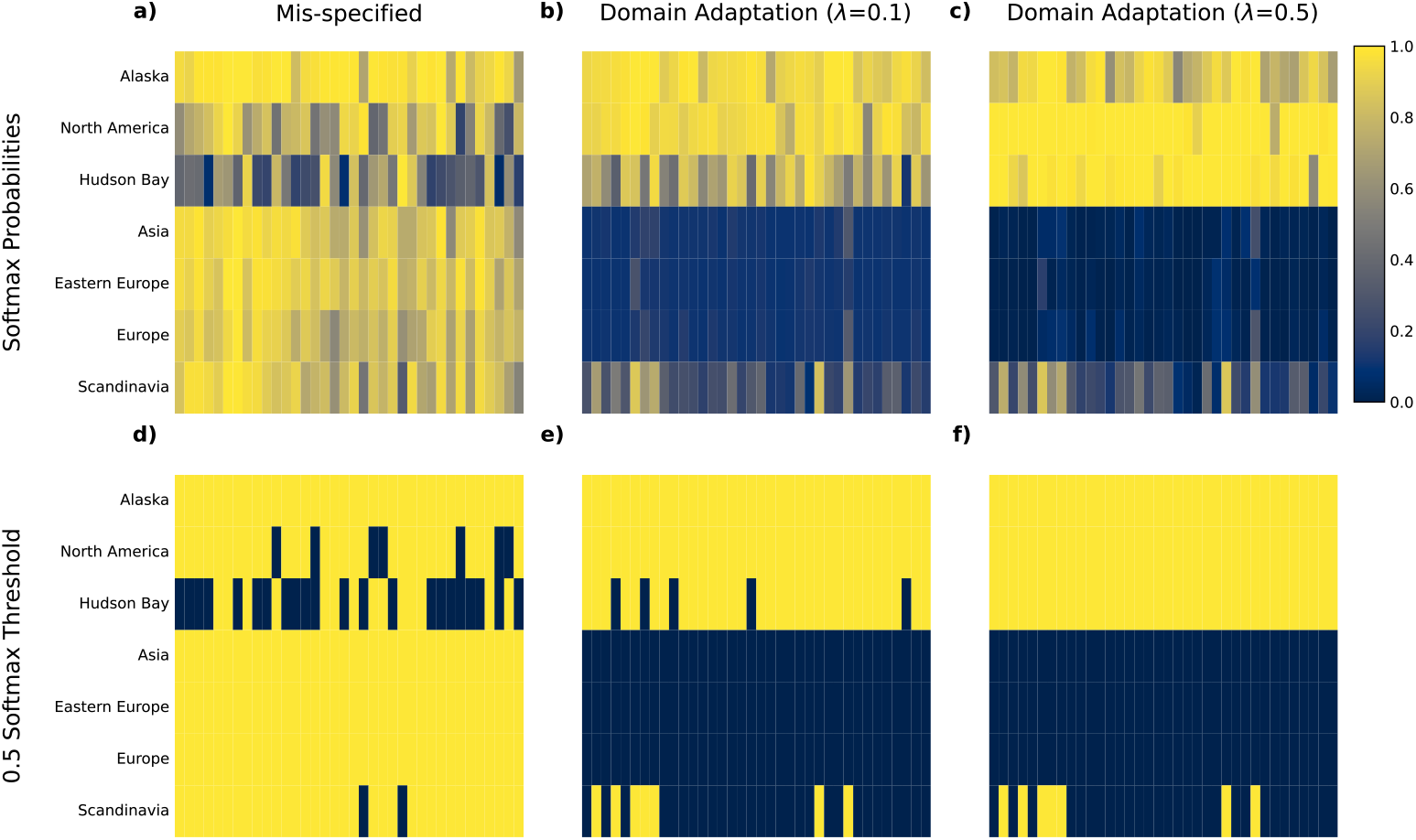
Results from tests of introgression between the ABC Island bears and other brown bear populations. Rows indicate the brown bear population, and columns correspond to the 36 chromosomes tested. (a-c) Average softmax probabilities across replicate networks with (a) *λ* = 0 (no domain adaptation), (b) *λ* = 0.1, and (c) *λ* = (d-f) A 50% threshold was applied to the softmax probabilities, such that a chromosome was only classified as introgressed if the average softmax probability of introgression across all replicates was > 0.5. Based on this threshold, chromosomes are classified as introgressed (yellow) or not introgressed (blue). Results are shown with (d) *λ* = 0 (no domain adaptation), (e) *λ* = 0.1, and (f) *λ* = 0.5.

Training networks with domain adaptation reduced support for introgression between geographically isolated populations of brown bears. When *λ* was set to 0.1, the mean softmax probabilities of introgression across replicates were much lower for most chromosomes in all populations, with a mean of 0.46 (SD = 0.46) (Fig. 8b). When *λ* was set to 0.5, the mean softmax probabilities of introgression across replicates were much lower for most chromosomes in all populations, with a mean of 0.46 (SD = 0.45) (Fig. 8c). Furthermore, when using domain adaptation the number of datasets classified as having introgression using a classification threshold of 0.5 was much lower (Fig. 8e-f). Notably, there are fewer tests supporting introgression between the ABC Island bears and geographically isolated populations (Fig. 8e-f), while introgression is inferred between the ABC Island bears and other bear populations from North America with greater frequency. Importance weight estimates vary across *λ* values as seen in the simulation studies, with the highest level of stability seen at *λ* = 0.1 (Fig. S12).

## Discussion

### The Promise of Domain Adaptation

Machine learning approaches used in population genetics typically rely on simulated training data due to the absence of real labeled datasets. As a consequence, the processes generating empirical data may not be modeled, and the simulated data used to train the machine learning model may differ substantially from empirical data. A mismatch between the training data distribution, referred to as the source domain, and the empirical data distribution, referred to as the target domain, can reduce the accuracy of inferences made with these tools (Wilson and Cook, 2020). We demonstrated the effectiveness of domain adaptation in overcoming this issue in inferences of introgression from population genetic data. Specifically, we tested the performance of CDAN, a type of adversarial domain adaptation (Long et al., 2018). With this approach, the network is trained to extract features shared across the source and target domains that can classify the data into the desired categories. In our case, the aim was to find features that could be used to classify the data as having past introgression between populations or not, even in the presence of unmodeled ghost introgression, a known cause of error in tests for introgression (Tricou et al., 2022; Beerli, 2004; Strasburg and Rieseberg, 2010).

We found that unmodeled ghost introgression negatively impacted inference using a standard CNN. However, using CDAN, we could detect introgression with very high accuracy, even in the presence of ghost introgression (Fig. 4). The CDAN was able to achieve this high level of accuracy using a relatively small amount of unlabeled data from the target domain, which bodes well for real-world applications of this approach. The much higher level of accuracy achieved with the CDAN indicates that it was able to effectively learn features informative for classification of the data that are shared between the source and target domains. Additionally, the features extracted by the CDAN exhibit a higher degree of similarity between the source and target domains compared to features extracted by a traditional CNN, demonstrating the alignment of the learned features extracted during training (Fig. 4c,f). When performing classification in the source domain, we did not find any evidence of a substantial negative impact of using domain adaptation, even though there is no domain shift in this case. This is promising, because it may not always be obvious that there is a domain shift in empirical systems, but the safer approach may be to use domain adaptation anyway. However, it remains likely that domain adaptation will not lead to the same level of accuracy expected in the absence of a domain shift, as was seen in Mo and Siepel (2023). Thus, if the user is aware of a particular domain shift, and the omitted process is computationally feasible to model, then it is likely preferable to train on the more appropriate domain than to use domain adaptation. However, it is not possible to know the true model *a priori*, and so one can never be sure that they are avoiding domain shifts. Based on our results, we recommend applying domain adaptation even in cases where there is no clear domain shift, and, at least, comparing results with and without domain adaptation.

Notably, we found that class imbalance could quickly degrade performance in the target domain. This led us to adopt importance-weighting, which attempts to accommodate imbalance in target data (Tachet des Combes et al., 2020). Using importance weights substantially improved performance on imbalanced data, except when all target datasets came from a single class (Fig. 5, Fig. 7). Thus, we caution against using CDAN on datasets for which the researcher suspects that all data come from a single class, and encourage the use of importance-weighting whenever implementing CDAN in evolutionary biology.

In general, we expect that domain adaptation should work well when there are features that are both informative for classification and shared across domains. Introgression has a clear and distinctive impact on the joint SFS, leading to increased allele sharing between populations, while ghost introgression skews the SFS, resulting in more low-frequency alleles unique to the recipient population (Fig. 1, Fig. S3, Fig. S4). While this skew was expected, some of the other patterns resulting from ghost introgression are more counterintuitive at first glance and only easily observed and interpreted when looking at the unnormalized SFS (Fig. S4). Notably, the depletion of alleles fixed in population 1 (not the recipient of ghost introgression) (Fig. 1, Fig. S3a, Fig. S4a) at first seems counterintuitive because ghost introgression should not impact allele frequencies in this population in the absence of migration between the focal populations. However, we are using the pairwise folded SFS. We suspect that some alleles were shared between population 1 and the ghost population, but absent in population 2 due to incomplete lineage sorting. In the presence of ghost introgression, such alleles can be introduced into population 2. The effects of this become more complex due to our use of the folded SFS rather than the polarized SFS. The introduction of these alleles into population 2 not only causes these alleles to no longer be absent in population 2 (which would move them down from the top right of the SFS), but also flips the minor and major alleles, resulting in a shift to the left quadrant of the SFS (more easily visualized without normalization; Fig. S4c). This occurs, also, in alleles not fixed, but segregating at intermediate-to-high frequencies in population 1. Thus, ghost introgression leads to a (very slight) excess of shared variants in the absence of migration. This may explain the greater excess of false negatives compared to false positives in the presence of ghost introgression Fig. 4). Despite this, both false negatives from the target data and false positives from the target data look rather distinct from true negatives and true positives, respectively (Fig. S13). Since ghost introgression clearly alters the SFS, but does not produce patterns that exactly match those produced by introgression between the focal populations, it is relatively straightforward for the network to learn features useful for classification but shared across domains. Applying domain adaptation would likely be less effective if the patterns produced by ghost introgression more closely matched those produced by introgression between the focal populations. Visualizing the SFS across all simulations (Fig. 1, Fig. S1) and across correctly and incorrectly classified datasets clearly indicates that this is not the case here (Fig. S13).

There is a clear parallel between domain shifts in machine learning and model misspecification in likelihood-based inference. Just as domain shifts lead to decreased performance in machine learning, model misspecifications, in which important processes are omitted from the model used for inference, lead to decreased performance in likelihood approaches. For example, previous work has demonstrated that selection misleads demographic inference (e.g., Smith and Hahn 2024; Johri et al. 2021; Schrider et al. 2016), that unaccounted for ghost introgression misleads inferences of migration (e.g., Beerli 2004), and that other common model violations negatively impact a variety of inferential tasks (e.g., Soni et al. 2024; Jensen et al. 2005). Attempts to ameliorate these impacts have, by necessity, tended to focus on a single model violation and either expand the model to include the relevant process, or use heuristic approaches to approximate the relevant process (e.g., Roux et al. 2016; Rougeux et al. 2017). Domain adaptation is unique in that, in theory, it can be used to address any domain shift and even multiple domain shifts at once. Furthermore, domain adaptation does not require knowledge of the causes of domain shifts. While further research is needed to explore the efficacy across domain shifts in more contexts, this may position machine learning to uniquely address the issue of model violations in evolutionary inference.

However, in the case explored here, ghost introgression did not negatively impact inference in fastsimcoal2 nearly as much as it impacted our machine learning approach. It remains to be seen in general whether machine learning or likelihood-based inference is more robust to model violations, and this problem warrants exploration across a wider range of conditions. It is plausible that machine learning algorithms are prone to over-fitting the training data, and thus more susceptible to model violations than other approaches. If this is the case, it is even more important to pair these approaches with mitigating strategies, like domain adaptation.

### Application to Empirical Data

When applied to empirical data from brown bears, the network trained without domain adaptation produced highly implausible results. We do not expect introgression to have occurred between the ABC Island bears and bears from Asia and Europe. The Bering land bridge that permitted the migration of brown bears into North America was inundated by rising sea levels by approximately 11,000 years ago, isolating North American brown bears from other brown bear populations (de Jong et al., 2023). The occurrence of gene flow between ABC Island brown bears and mainland brown bears in North America, on the other hand, has been previously demonstrated (Cahill et al., 2013). Despite this expectation, the original network inferred introgression between Asian and European brown bears and the ABC Island bears across many chromosomes (Fig. 8a,d). Using domain adaptation reduced support for introgression, particularly between geographically isolated populations (Fig. 8b-c,e-f), and we suspect that these results are far more reliable than those without domain adaptation. There was one notable exception. Even with domain adaptation, we found some support for introgression between the ABC Island bears and bears from Scandinavia (Fig. 8). While we view this as biologically improbable, it is possible that this particular classification is plagued by some other domain shift not ameliorated by our current classifiers (see below).

### Challenges and Future Directions

Several questions remain, particularly concerning the application of our CDAN to empirical data. Below, we discuss additional sources of misspecification and the use of summary statistics in more detail. In our simulations, there was a single source of misspecification–ghost introgression. However, myriad other sources of misspecification may impact empirical datasets. For example, in addition to ghost introgression, background selection has been shown to negatively impact tests for introgression (Smith and Hahn, 2024). Additional sources of misspecification could include genotyping error, ghost introgression from multiple populations, population size changes, misspecified mutation and recombination rates, or unmodelled variation in recombination and mutation rates. Previous work has shown mixed impacts of such misspecifications, with misspecified recombination rates having minimal impacts on demographic inference in a deep learning approach (Sheehan and Song, 2016), unmodelled variation in mutation and recombination rates negatively impacting population size inference in approximate Likelihood approaches (Soni et al., 2024), and minimal impacts of misspecified demography on inferences of introgression in a machine learning approach (Schrider et al., 2018). Fortunately, our domain adaptation approach should be agnostic to the source of misspecification and should work as long as features exist that are shared across source and target domains which can be used for classification. However, there may be sources of model misspecification that alter the data in ways that domain adaptation cannot overcome. We also simulated a fairly modest level of misspecification, but it has been shown that domain adaptation is sensitive to the degree of misspecification (Mo and Siepel, 2023). Future work should explore different sources of model misspecification, interactions between sources of model misspecification, and the limits of domain adaptation in accommodating model misspecification.

Here, we used a summary statistic, the site frequency spectrum, as input into the network. In our tests using simulated data, this resulted in near-perfect accuracy. However, using a single summary statistic will result in a failure to capture all of the signal from genetic data, which may become a problem when models and inferential goals become more complex. Future work should evaluate the efficacy of domain adaptation when using additional summary statistics to detect introgression (e.g., those used in Schrider et al. 2018), including statistics related to linkage disequilibrium. One of the exciting possibilities of using CNNs for analyzing genetic data is that the raw DNA alignment can be used as input, which results in less loss of information as compared to the use of summary statistics (Flagel et al., 2019). Analyzing alignments directly could also facilitate domain adaptation by increasing the variety of features that could be learned for classification, which could in turn broaden the number of features available for classification and shared across domains. However, CNNs using raw sequence alignments are more challenging to implement as they come with greater computational burden, and trained models cannot be as readily re-used for datasets with different alignment lengths. Using combinations of several different summary statistics as input may yield improvements in domain adaptation while remaining flexible and easier to implement (Schrider et al., 2018), and future work should explore these options.

## Conclusions

Our results demonstrate both the fragility of traditional machine learning approaches in the presence of domain shifts and the effectiveness of domain adaptation in population genetics. The high error rates associated with a relatively mild (and likely common) model misspecification warrant caution when applying supervised machine learning approaches to population genetic data. Avoiding model misspecification is impossible, as all models are necessarily simplifications of reality, and our results suggest that standard supervised learning approaches may not be robust to common model violations. Despite these concerning results, our results also suggest a promising path forward. The domain adaptation approach applied here is computationally tractable, requires no labeled data from the target domain, and apparently does not decrease accuracy even in the absence of misspecification. We are cautiously optimistic, and suggest that future machine learning approaches in population genetics should strongly consider the use of unsupervised domain adaptation.

## Code Availability

All code used for this project is available at: https://doi.org/10.5281/zenodo.22802059

## Acknowledgements

Thank you to members of the Smith Lab for providing feedback on this manuscript.

