## Supplemental Figures for "The reasonable effectiveness of domain adaptation for inference of introgression"

Title: The reasonable effectiveness of domain adaptation for inference of introgression
Authors: Kerry A. Cobb[1,*], Megan L. Smith[1]

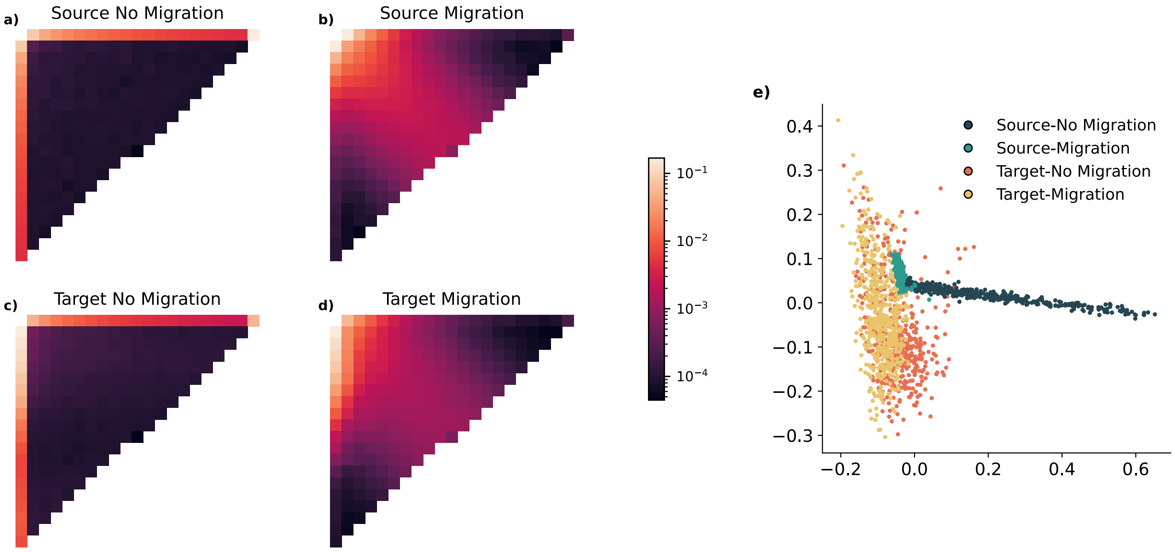


Figure S1. SFS generated under the general simulation study. (a-d) The average normalized SFS for test data simulated under the general scenario. e) PCA of the average normalized SFS for test data simulated under the general scenario.

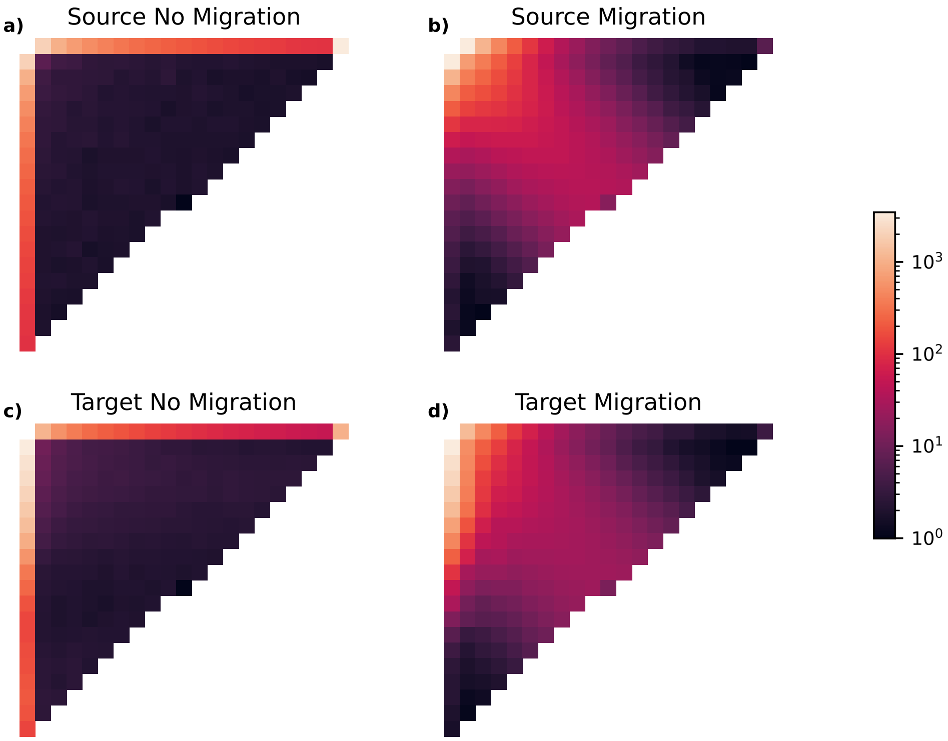


Figure S2. Unnormalized SFS generated under the general simulation study. (a-d) The average unnormalized SFS for test data simulated under the general scenario.

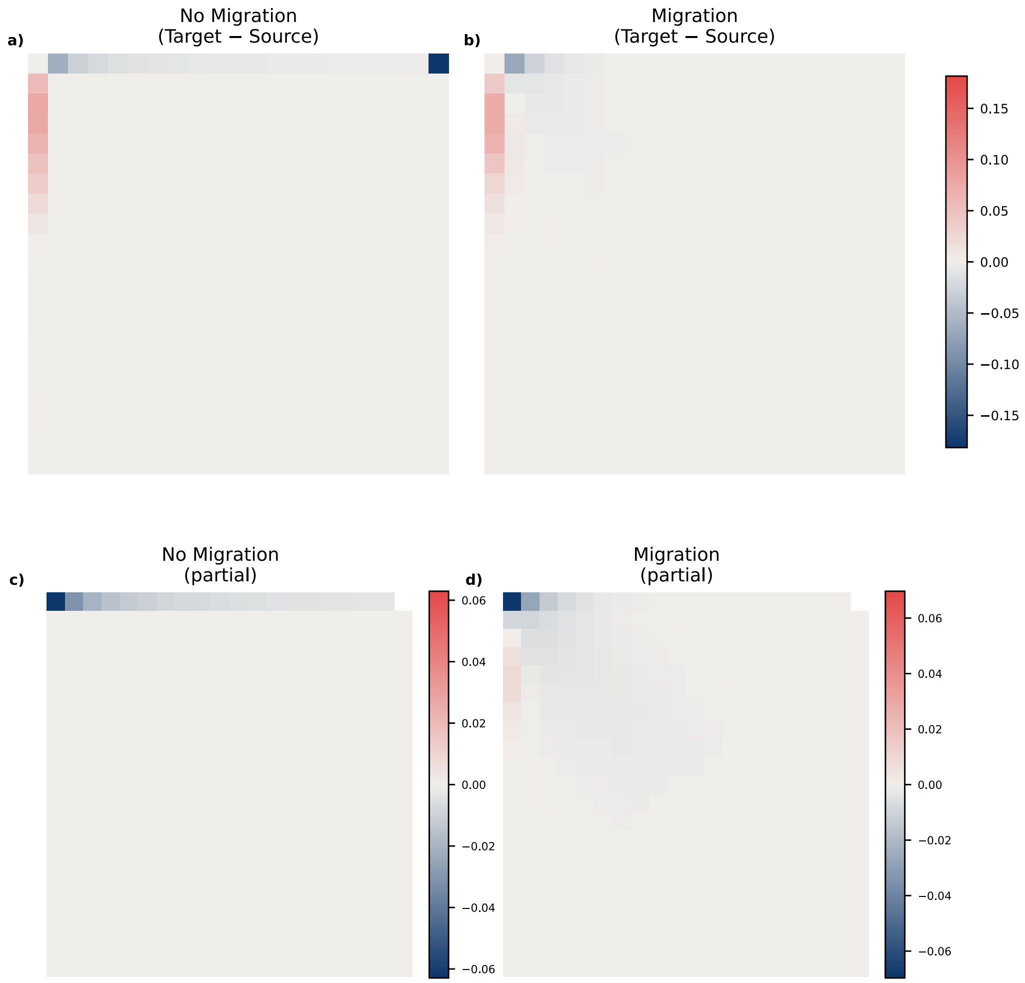


Figure S3. The difference between normalized SFS generated under source and target scenarios in the general simulation scenario. a) The difference in the absence of migration; b) The difference in the presence of migration; c) The difference in the absence of migration excluding the first column and the final cell of the first row to facilitate visualization. d) The difference in the presence of migration excluding the first column and the final cell of the first row to facilitate visualization.

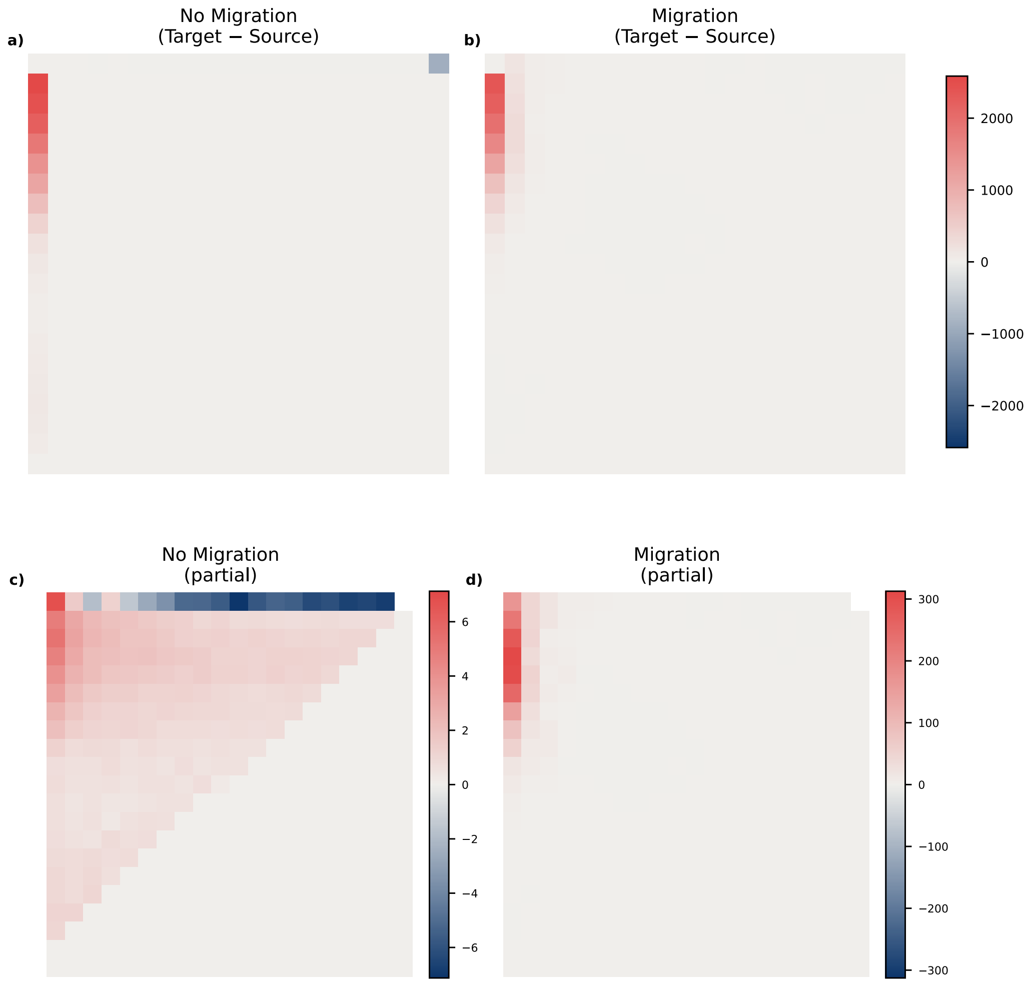


Figure S4. The difference between unnormalized SFS generated under source and target scenarios in the general simulation scenario. a) The difference in the absence of migration; b) The difference in the presence of migration; c) The difference in the absence of migration excluding the first column and the final cell of the first row to facilitate visualization. d) The difference in the presence of migration excluding the first column and the final cell of the first row to facilitate visualization.

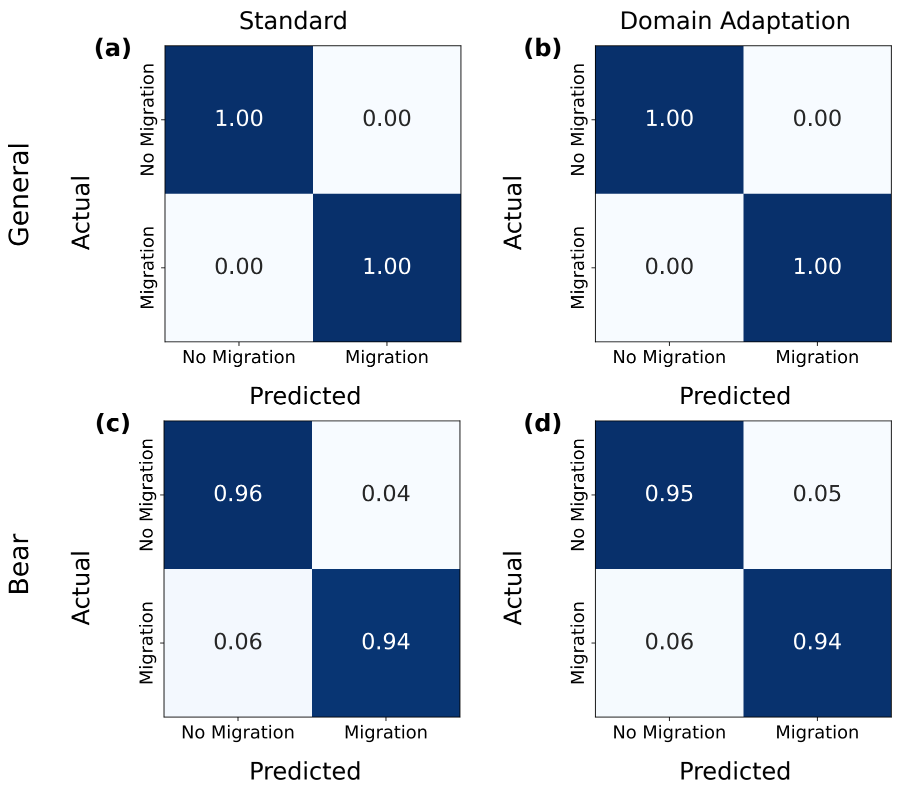


Figure S5. Results from datasets generated under the source domain (i.e., without ghost introgression). (a-b) Results under the general simulation scenario. Confusion matrices show the proportion of test replicates in each category without (a) and with (d) domain adaptation. (c-d) Results under the bear simulation scenario. Confusion matrices show the proportion of test replicates in each category without (c) and with (d) domain adaptation. Values were averaged across the ten replicate networks. All values are rounded down to the nearest hundredths position. See Table S1 for original values.

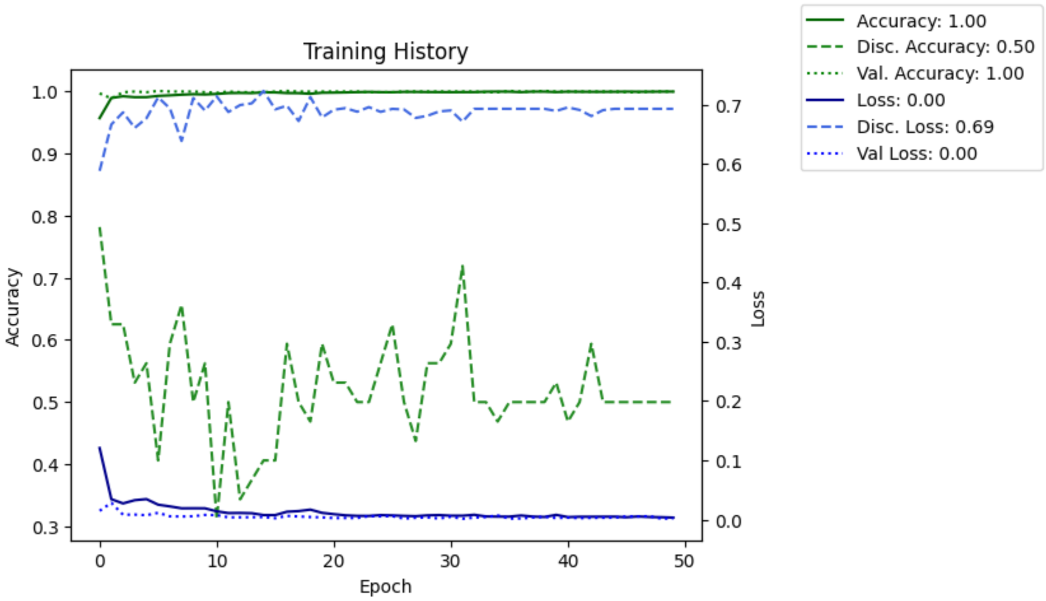


Figure S6. Training accuracy and loss of the classifier (solid lines) and discriminator (dashed lines) at each training epoch for one representative training replicate using general scenario training data.

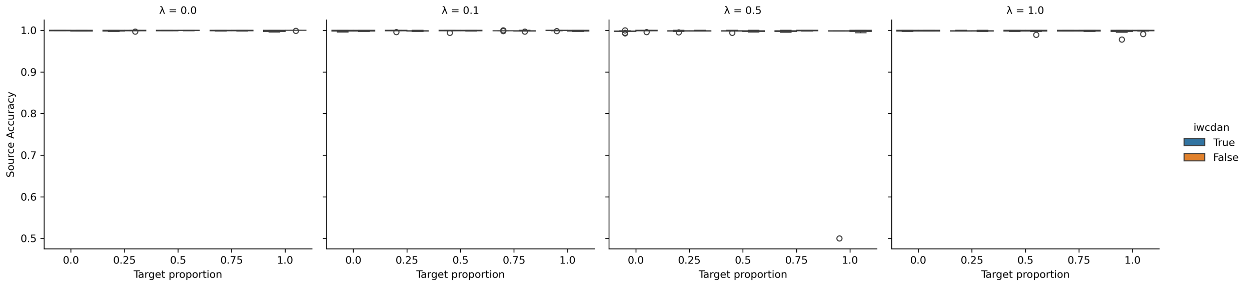


Figure S7. Results across imbalanced datasets under the general simulation scenario under the source domain. The x-axis is the proportion of the 50 target datasets coming from the no migration class. Blue boxplots show results with importance weighting, and orange boxplots show results with importance weighting.

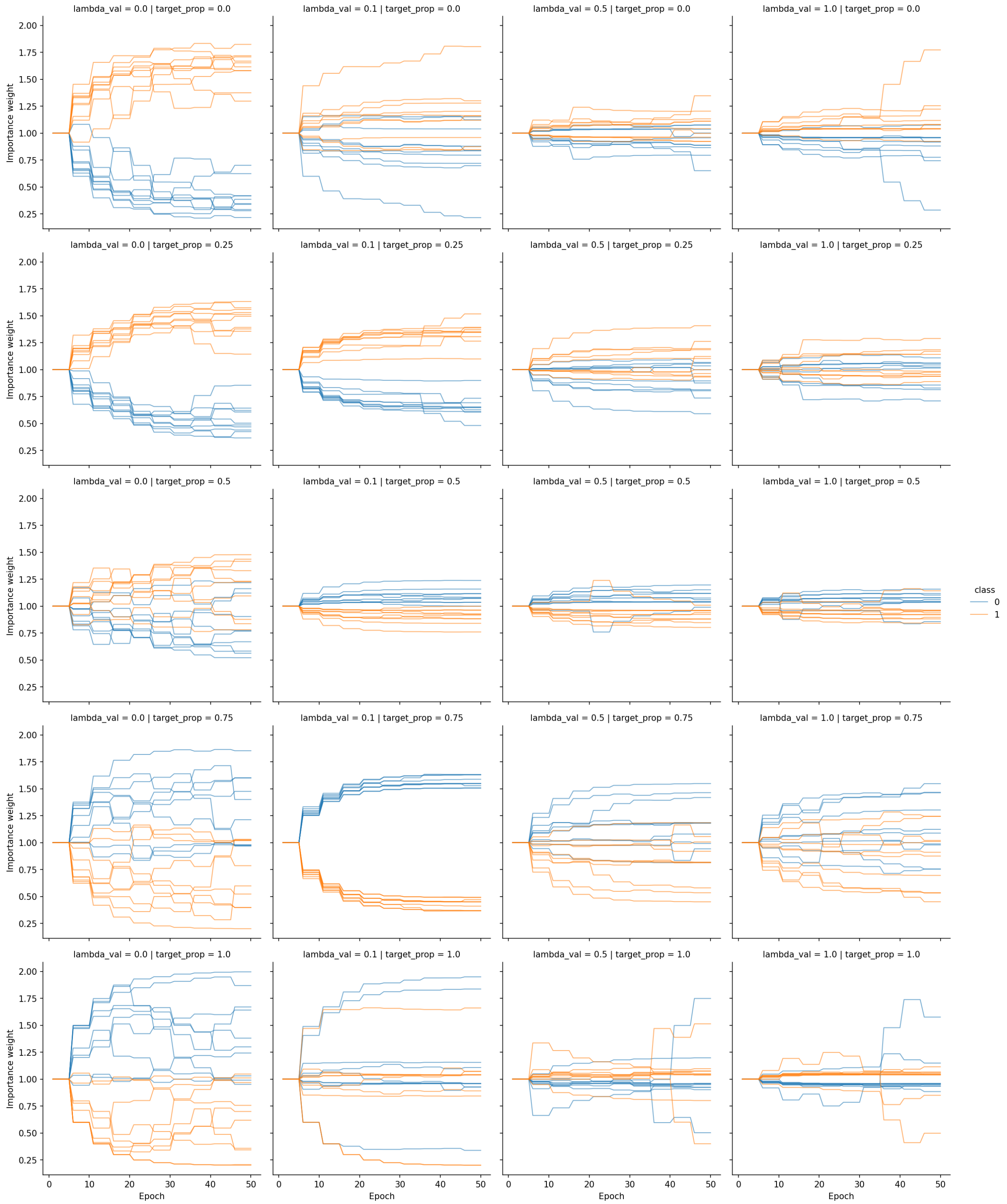


Figure S8. Importance weight estimates across epochs for different values of $\lambda$ and the target proportion for the general simulation scenario.

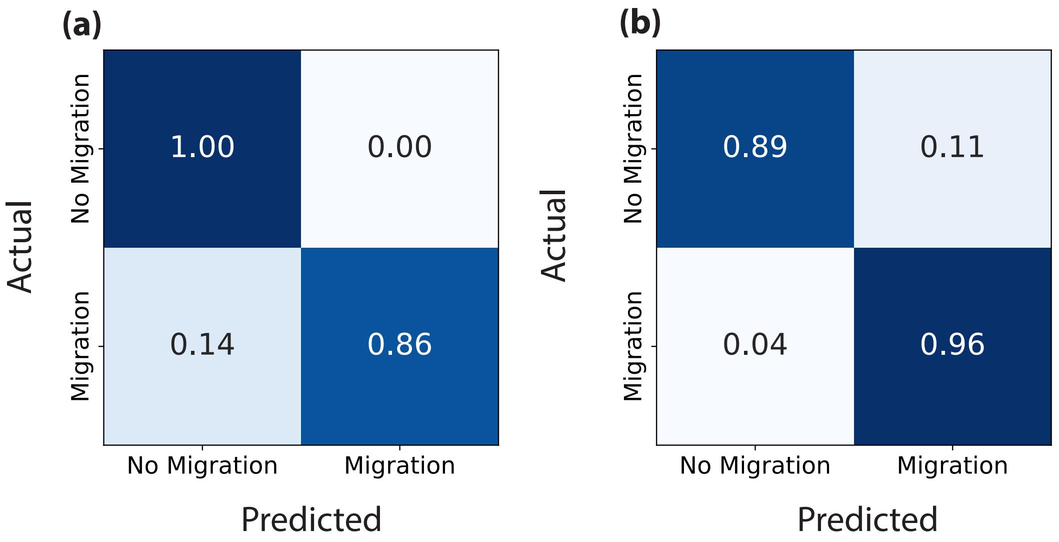


Figure S9. Results under the bear simulation on datasets generated under the source (a) and target (b) domains. Confusion matrices show the proportion of test replicates in each category without domain adaptation at a threshold of 0.9.

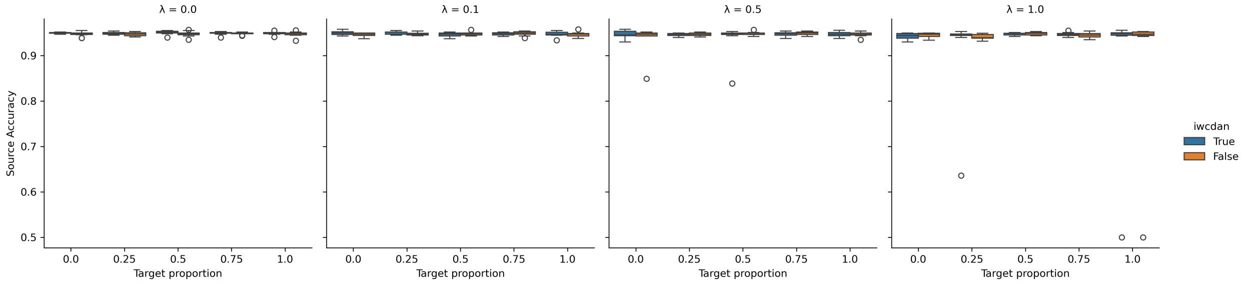


Figure S10. Results across imbalanced datasets under the bear simulation scenario under the source domain. The x-axis is the proportion of the 50 target datasets coming from the no migration class. Blue boxplots show results with importance weighting, and orange boxplots show results with importance weighting.

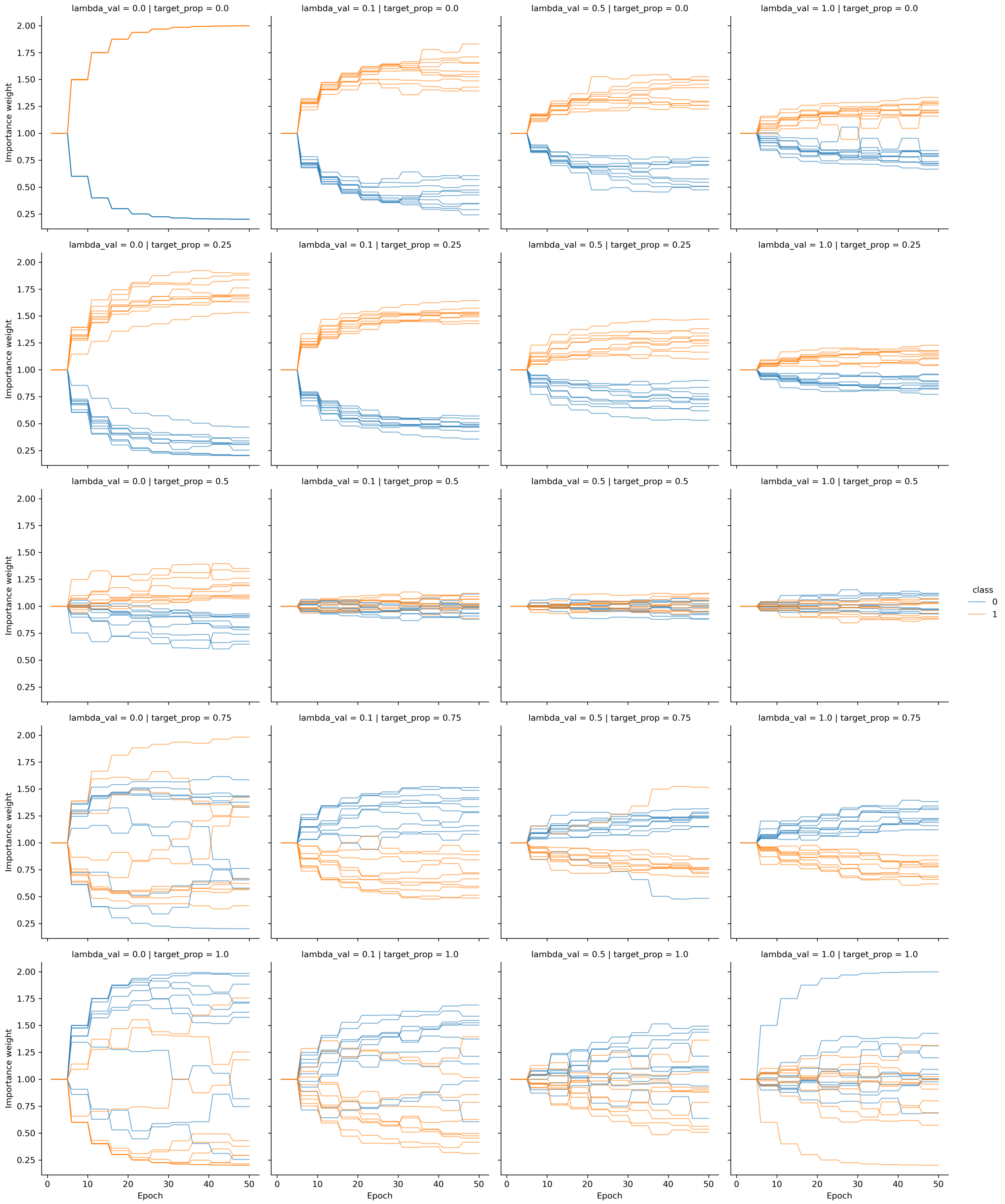


Figure S11. Importance weight estimates across epochs for different values of $\lambda$ and the target proportion for the bear simulation scenario.

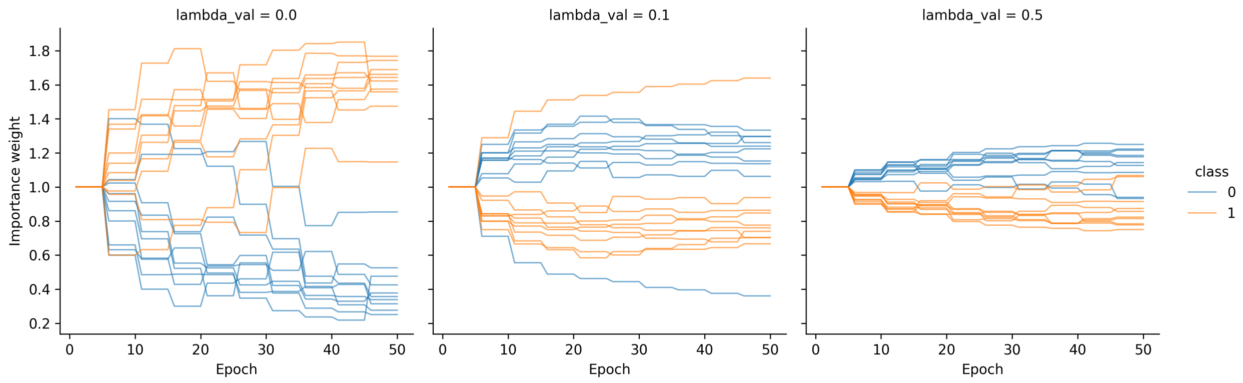


Figure S12. Importance weight estimates across epochs for different values of $\lambda$ and for the empirical data.

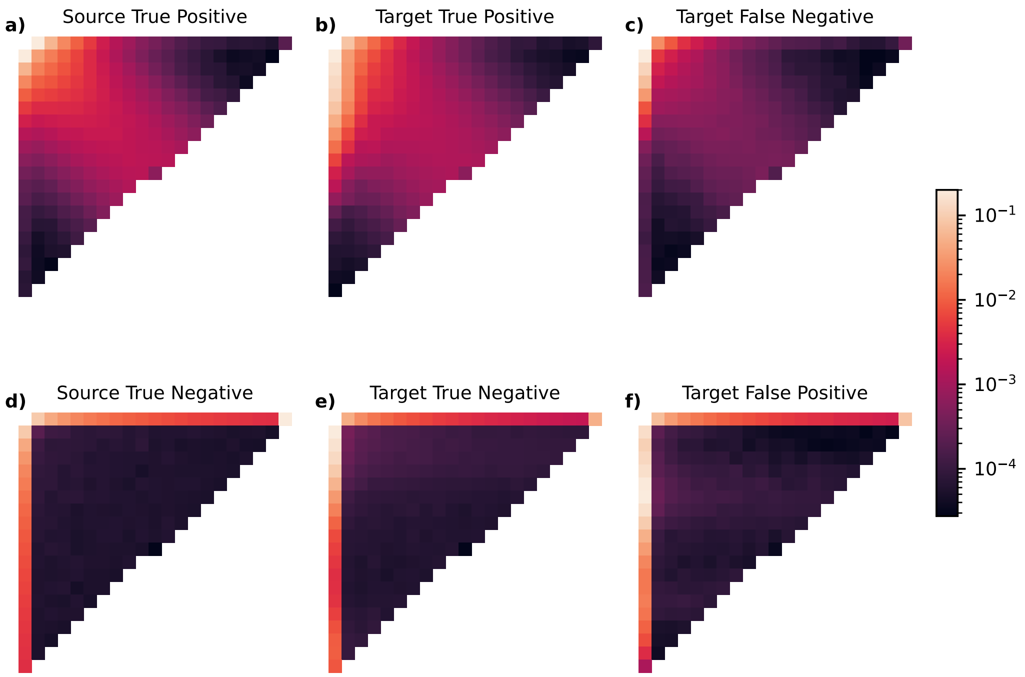


Figure S13. Average normalized site frequency spectrum (SFS) for test data simulated under the general scenario. Values of each cell are the mean of all SFS for a given error category. The recipient population of ghost introgression in target domain datasets is on the row axis.

Table S1. Unrounded mean error rates across tests of trained network replicates.

| Scenario | $\lambda$ | Domain | False Positive | False Negative | AUC |
|---|---|---|---|---|---|
| General | 0.0000 | Source | 0.0 | 0.0014 | 0.9999916 |
| General | 0.0 | Target | 0.2810 | 0.1798 | 0.8615885999999999 |
| General | 1.0 | Source | 0.0002 | 0.0030 | 0.9999972 |
| General | 1.0 | Target | 0.0086 | 0.0504 | 0.994225 |
| Bear | 0.0 | Source | 0.0414 | 0.0622 | 0.990888 |
| Bear | 0.0 | Target | 0.1760 | 0.0208 | 0.9831799999999999 |
| Bear | 0.5 | Source | 0.0470 | 0.0568 | 0.9904892 |
| Bear | 0.5 | Target | 0.0438 | 0.0482 | 0.988615 |

Table S2: Bear samples used in our empirical tests for introgression.

| BioSample ID | SRA Run ID | Population | Publication | Lat | Lon |
|---|---|---|---|---|---|
| SAMEA4762870 | ERR2678638 | Europe | Barlow et al. 2018 | 46.445 | 14.01778 |
| SAMN07422262 | SRR5878348 | Europe | Benazo et al. 2017 | 42.03908 | 13.43847 |
| SAMN07422262 | SRR5878360 | Europe | Benazo et al. 2017 | 42.03908 | 13.43847 |
| SAMN07422269 | SRR5878338 | Europe | Benazo et al. 2017 | 48.14816 | 17.10674 |
| SAMN07422269 | SRR5878353 | Europe | Benazo et al. 2017 | 48.14816 | 17.10674 |
| SAMN07422272 | SRR5878341 | Europe | Benazo et al. 2017 | 46.15124 | 14.99546 |
| SAMN07422272 | SRR5878346 | Europe | Benazo et al. 2017 | 46.15124 | 14.99546 |
| SAMN07422268 | SRR5878351 | Europe | Benazo et al. 2018 | 39.07421 | 21.82431 |
| SAMN07422268 | SRR5878354 | Europe | Benazo et al. 2018 | 39.07421 | 21.82431 |
| SAMN02045559 | SRR830337 | Alaska | Cahill et al. 2013 | 63.12989 | -151.197 |
| SAMN02045560 | SRR830213 | ABC Islands | Cahill et al. 2015 | 57.44 | -134.2 |
| SAMN03247209 | SRR1692419 | ABC Islands | Cahill et al. 2015 | 57.87083 | -135.772 |
| SAMN02256316 | SRR935596 | ABC Islands | Liu et al. 2014 | 56.9508 | -134.942 |
| SAMN02256316 | SRR935603 | ABC Islands | Liu et al. 2014 | 56.9508 | -134.942 |
| SAMN02256316 | SRR935610 | ABC Islands | Liu et al. 2014 | 56.9508 | -134.942 |
| SAMN02256316 | SRR935618 | ABC Islands | Liu et al. 2014 | 56.9508 | -134.942 |
| SAMN02256316 | SRR941809 | ABC Islands | Liu et al. 2014 | 56.9508 | -134.942 |
| SAMN02256316 | SRR941812 | ABC Islands | Liu et al. 2014 | 56.9508 | -134.942 |
| SAMN02256317 | SRR935597 | ABC Islands | Liu et al. 2014 | 56.9508 | -134.942 |
| SAMN02256317 | SRR935604 | ABC Islands | Liu et al. 2014 | 56.9508 | -134.942 |
| SAMN02256317 | SRR935611 | ABC Islands | Liu et al. 2014 | 56.9508 | -134.942 |
| SAMN02256317 | SRR935620 | ABC Islands | Liu et al. 2014 | 56.9508 | -134.942 |
| SAMN02256317 | SRR935629 | ABC Islands | Liu et al. 2014 | 56.9508 | -134.942 |
| SAMN02256318 | SRR935598 | ABC Islands | Liu et al. 2014 | 57.87083 | -135.772 |
| SAMN02256318 | SRR935605 | ABC Islands | Liu et al. 2014 | 57.87083 | -135.772 |
| SAMN02256318 | SRR935612 | ABC Islands | Liu et al. 2014 | 57.87083 | -135.772 |
| SAMN02256318 | SRR935621 | ABC Islands | Liu et al. 2014 | 57.87083 | -135.772 |
| SAMN02256318 | SRR941786 | ABC Islands | Liu et al. 2014 | 57.87083 | -135.772 |
| SAMN02256320 | SRR935600 | ABC Islands | Liu et al. 2014 | 57.87083 | -135.772 |
| SAMN02256320 | SRR935607 | ABC Islands | Liu et al. 2014 | 57.87083 | -135.772 |
| SAMN02256320 | SRR935614 | ABC Islands | Liu et al. 2014 | 57.87083 | -135.772 |
| SAMN02256320 | SRR935623 | ABC Islands | Liu et al. 2014 | 57.87083 | -135.772 |
| SAMN02256320 | SRR941808 | ABC Islands | Liu et al. 2014 | 57.87083 | -135.772 |
| SAMN02256321 | SRR935601 | ABC Islands | Liu et al. 2014 | 57.73333 | -134.333 |
| SAMN02256321 | SRR935608 | ABC Islands | Liu et al. 2014 | 57.73333 | -134.333 |
| SAMN02256321 | SRR935615 | ABC Islands | Liu et al. 2014 | 57.73333 | -134.333 |
| SAMN02256321 | SRR935619 | ABC Islands | Liu et al. 2014 | 57.73333 | -134.333 |
| SAMN02256321 | SRR941810 | ABC Islands | Liu et al. 2014 | 57.73333 | -134.333 |
| SAMN02256321 | SRR941813 | ABC Islands | Liu et al. 2014 | 57.73333 | -134.333 |
| SAMN02256315 | SRR935592 | Eastern Europe | Liu et al. 2014 | 61.28333 | 28.83333 |
| SAMN02256315 | SRR935595 | Eastern Europe | Liu et al. 2014 | 61.28333 | 28.83333 |
| SAMN02256315 | SRR935624 | Eastern Europe | Liu et al. 2014 | 61.28333 | 28.83333 |

Continued on next page

Table S2 – continued from previous page

| BioSample ID | SRA Run ID | Population | Publication | Lat | Lon |
|---|---|---|---|---|---|
| SAMN02256315 | SRR935628 | Eastern Europe | Liu et al. 2014 | 61.28333 | 28.83333 |
| SAMN02256322 | SRR935602 | North America | Liu et al. 2014 | 46.96526 | -109.534 |
| SAMN02256322 | SRR935609 | North America | Liu et al. 2014 | 46.96526 | -109.534 |
| SAMN02256322 | SRR935616 | North America | Liu et al. 2014 | 46.96526 | -109.534 |
| SAMN02256322 | SRR935617 | North America | Liu et al. 2014 | 46.96526 | -109.534 |
| SAMN02256322 | SRR941811 | North America | Liu et al. 2014 | 46.96526 | -109.534 |
| SAMN02256322 | SRR941814 | North America | Liu et al. 2014 | 46.96526 | -109.534 |
| SAMN02256314 | SRR935591 | Scandinavia | Liu et al. 2014 | 66.60665 | 19.82324 |
| SAMN02256314 | SRR935625 | Scandinavia | Liu et al. 2014 | 66.60665 | 19.82324 |
| SAMN02256314 | SRR935627 | Scandinavia | Liu et al. 2014 | 66.60665 | 19.82324 |
| SAMN01057688 | SRR518710 | ABC Islands | Miller et al. 2012 | 57.44 | -134.2 |
| SAMN01057688 | SRR518711 | ABC Islands | Miller et al. 2012 | 57.44 | -134.2 |
| SAMN01057690 | SRR518712 | Alaska | Miller et al. 2012 | 60.55444 | -151.258 |
| SAMN01057690 | SRR518713 | Alaska | Miller et al. 2012 | 60.55444 | -151.258 |
| SAMN09907428 | SRR7758718 | North America | Taylor et al. 2018 | 58.747 | -128.878 |
| SAMN32301304 | SRR22801890 | ABC Islands | de Jong et al. 2023 | 57.73333 | -134.333 |
| SAMN32301305 | SRR22801879 | ABC Islands | de Jong et al. 2023 | 57.73333 | -134.333 |
| SAMN32301308 | SRR22801818 | Alaska | de Jong et al. 2023 | 68.143 | -151.736 |
| SAMN32301309 | SRR22801807 | Alaska | de Jong et al. 2023 | 68.143 | -151.736 |
| SAMN32301310 | SRR22801856 | Alaska | de Jong et al. 2023 | 68.143 | -151.736 |
| SAMN32301311 | SRR22801845 | Alaska | de Jong et al. 2023 | 68.143 | -151.736 |
| SAMN32301394 | SRR22801842 | Alaska | de Jong et al. 2023 | 55.81 | -166.66 |
| SAMN32301395 | SRR22801841 | Alaska | de Jong et al. 2023 | 55.81 | -166.66 |
| SAMN32301396 | SRR22801840 | Alaska | de Jong et al. 2023 | 66.94 | -160.6 |
| SAMN32301397 | SRR22801839 | Alaska | de Jong et al. 2023 | 70.29 | -148.79 |
| SAMN32301312 | SRR22801900 | Asia | de Jong et al. 2023 | 52.43055 | 140.3289 |
| SAMN32301313 | SRR22801899 | Asia | de Jong et al. 2023 | 46.0321 | 136.677 |
| SAMN32301314 | SRR22801898 | Asia | de Jong et al. 2023 | 45.4562 | 137.185 |
| SAMN32301316 | SRR22801896 | Asia | de Jong et al. 2023 | 57.645 | 136.23 |
| SAMN32301336 | SRR22801874 | Asia | de Jong et al. 2023 | 59.5638 | 150.8035 |
| SAMN32301339 | SRR22801871 | Asia | de Jong et al. 2023 | 59.5638 | 150.8035 |
| SAMN32301340 | SRR22801870 | Asia | de Jong et al. 2023 | 56.65 | 124.7 |
| SAMN32301342 | SRR22801867 | Asia | de Jong et al. 2023 | 60.02505 | 123.3926 |
| SAMN32301343 | SRR22801838 | Asia | de Jong et al. 2023 | 64.16806 | 116.4672 |
| SAMN32301345 | SRR22801836 | Asia | de Jong et al. 2023 | 59.6 | 153.15 |
| SAMN32301334 | SRR22801876 | Eastern Europe | de Jong et al. 2023 | 59.008 | 26.226 |
| SAMN32301335 | SRR22801875 | Eastern Europe | de Jong et al. 2023 | 59.2672 | 27.552 |
| SAMN32301346 | SRR22801835 | Eastern Europe | de Jong et al. 2023 | 61.6934 | 29.729 |
| SAMN32301347 | SRR22801834 | Eastern Europe | de Jong et al. 2023 | 61.91971 | 27.81635 |
| SAMN32301381 | SRR22801857 | Eastern Europe | de Jong et al. 2023 | 58.0 | 56.31667 |
| SAMN32301382 | SRR22801855 | Eastern Europe | de Jong et al. 2023 | 58.0 | 56.31667 |
| SAMN32301383 | SRR22801854 | Eastern Europe | de Jong et al. 2023 | 58.0 | 56.31667 |
| SAMN32301384 | SRR22801853 | Eastern Europe | de Jong et al. 2023 | 58.0 | 56.31667 |
| SAMN32301385 | SRR22801852 | Eastern Europe | de Jong et al. 2023 | 60.3262 | 56.423 |
| SAMN32301366 | SRR22801813 | Europe | de Jong et al. 2023 | 44.43996 | 26.09631 |
| SAMN32301367 | SRR22801812 | Europe | de Jong et al. 2023 | 44.43996 | 26.09631 |
| SAMN32301368 | SRR22801811 | Europe | de Jong et al. 2023 | 44.43996 | 26.09631 |
| SAMN32301369 | SRR22801810 | Europe | de Jong et al. 2023 | 44.43996 | 26.09631 |
| SAMN32301370 | SRR22801809 | Europe | de Jong et al. 2023 | 44.43996 | 26.09631 |
| SAMN32301302 | SRR22801902 | North America | de Jong et al. 2023 | 57.98417 | -127.788 |
| SAMN32301303 | SRR22801901 | North America | de Jong et al. 2023 | 57.98417 | -127.788 |
| SAMN32301317 | SRR22801895 | North America | de Jong et al. 2023 | 49.93 | -117.627 |
| SAMN32301387 | SRR22801850 | North America | de Jong et al. 2023 | 44.0682 | -114.742 |
| SAMN32301388 | SRR22801849 | North America | de Jong et al. 2023 | 48.1978 | -114.316 |
| SAMN32301389 | SRR22801848 | North America | de Jong et al. 2023 | 44.03971 | -109.541 |
| SAMN32301390 | SRR22801847 | North America | de Jong et al. 2023 | 48.8 | -117.255 |

Continued on next page

Table S2 – continued from previous page

| BioSample ID | SRA Run ID | Population | Publication | Lat | Lon |
|---|---|---|---|---|---|
| SAMN32301391 | SRR22801846 | North America | de Jong et al. 2023 | 44.56 | -111.444 |
| SAMN32301358 | SRR22801822 | Scandinavia | de Jong et al. 2023 | 69.96887 | 23.27165 |
| SAMN32301360 | SRR22801820 | Scandinavia | de Jong et al. 2023 | 69.4439 | 25.80482 |
| SAMN32301361 | SRR22801819 | Scandinavia | de Jong et al. 2023 | 69.4439 | 25.80482 |
| SAMN32301373 | SRR22801865 | Scandinavia | de Jong et al. 2023 | 60.83333 | 11.6666 |
| SAMN32301374 | SRR22801864 | Scandinavia | de Jong et al. 2023 | 60.83333 | 11.6666 |
| SAMN32301376 | SRR22801862 | Scandinavia | de Jong et al. 2023 | 63.16666 | 10.3333 |
| SAMN32301377 | SRR22801861 | Scandinavia | de Jong et al. 2023 | 63.16666 | 10.3333 |
| SAMN32301378 | SRR22801860 | Scandinavia | de Jong et al. 2023 | 63.16666 | 10.3333 |
| SAMN32301379 | SRR22801859 | Scandinavia | de Jong et al. 2023 | 63.16666 | 10.3333 |
